# The rodent vaginal microbiome across the estrous cycle and the effect of genital nerve electrical stimulation

**DOI:** 10.1101/647545

**Authors:** Micah Levy, Christine M. Bassis, Eric Kennedy, Katie E. Yoest, Jill B. Becker, Jason Bell, Mitchell B. Berger, Tim M. Bruns

**Author notes:** Corresponding author (TMB).

## Abstract

Treatment options are limited for the approximately 40% of postmenopausal women worldwide who suffer from female sexual dysfunction (FSD). Neural stimulation has shown potential as a treatment for genital arousal FSD, however the mechanisms for its improvement are unknown. One potential cause of some cases of genital arousal FSD are changes to the composition of the vaginal microbiota, which is associated with vulvovaginal atrophy. The primary hypothesis of this study was that neural stimulation may induce healthy changes in the vaginal microbiome, thereby improving genital arousal FSD symptoms. In this study we used healthy rats, which are a common animal model for sexual function, however the rat vaginal microbiome is understudied. Thus this study also sought to examine the composition of the rat vaginal microbiota. Treatment rats (n=5) received 30 minutes of cutaneous electrical stimulation targeting the genital branch of the pudendal nerve, and Control animals (n=4) had 30-minute sessions without stimulation. Vaginal lavage samples were taken during a 14-day baseline period including multiple estrous periods and after twice-weekly 30-minute sessions across a six-week trial period. Analysis of 16S rRNA gene sequences was used to characterize the rat vaginal microbiota in baseline samples and determine the effect of stimulation. We found that the rat vaginal microbiota is dominated by *Proteobacteria, Firmicutes*, and *Actinobacteria*, which changed in relative abundance during the estrous cycle and in relationship to each other. While the overall stimulation effects were unclear in these healthy rats, some Treatment animals had less alteration in microbiota composition between sequential samples than Control animals, suggesting that stimulation may help stabilize the vaginal microbiome. Future studies may consider additional physiological parameters, in addition to the microbiome composition, to further examine vaginal health and the effects of stimulation.

## Introduction

Female sexual dysfunction (FSD) affects up to 41% of postmenopausal women worldwide [1], and between 40-50% of all women report experiencing at least one symptom of FSD [2]. Between 16-25% of women report having difficulty orgasming, and between 8-28% of women report having low genital arousal and poor lubrication [3], [4]. FSD is inversely related to quality of life both physically and psychologically [2], [5]. Genital arousal FSD has been associated with Vulvovaginal Atrophy (VVA) in some postmenopausal women [6]. Symptoms of VVA include vaginal dryness, itching, irritation, and dyspareunia [6]. FSD and VVA stemming from postmenopausal hypoestrogenism induces an elevation of the vaginal pH [7]. High pH levels are associated with reductions of *Lactobacillus* species and an increase in the growth of other bacterial species such as group B *Streptococcus, Staphylococcus*, coliforms, and diphtheroids [7]. This reduction of a dominating species and overall increase in the diversity of the vaginal microbiota is associated with an overall reduction in vaginal health in postmenopausal women [8]. Postmenopausal women are 7.8 times more likely to have a more diverse vaginal microbiota than are premenopausal women [9], and this may be due to decreases in their concentrations of ovarian hormones [10]. This suggests that hormone changes associated with menopause may have an effect on the vaginal microbiome.

Though studies suggest that female sexual dysfunction is more prevalent than male sexual dysfunction [11], treatment options for FSD and VVA are limited compared to treatment for male sexual dysfunction. Current treatment options for symptoms of VVA include non-hormonal lubrication and estrogen treatments [7], though hormone therapy is not always recommended for long term dysfunction [12]. Various pharmaceutical treatments have been evaluated. While Sildenafil has shown improvements in clitoral blood flow and sexual arousal in postmenopausal women [13], its clinical efficacy and high risk of adverse effects such as headaches, flushing, rhinitis, and nausea have led to concern for its true clinical benefit in women [14], [15]. Flibanserin can improve sexual desire in postmenopausal women [16], however its efficacy in improving the physiological problems associated with FSD have not been demonstrated [17], [18].

Recent studies have shown that peripheral nerve stimulation, a third-line treatment for bladder dysfunction, can improve FSD symptoms. The implant of a sacral neuromodulation stimulator, which targets a sacral spinal nerve that contains proximal pudendal nerve fibers, for incontinence and overactive bladder has had positive benefits for FSD [19]. Weekly posterior tibial nerve stimulation (PTNS) sessions with a percutaneous needle electrode for similar patients has also shown promise [20], [21]. Recently we showed that 30-minute weekly sessions of skin-surface PTNS or stimulation on either side of the clitoris to target the genital branch of the pudendal nerve in women without bladder problems could improve the genital arousal aspects of FSD [22]. The underlying mechanisms of these improvements are not known. In preclinical studies, we found that increases in rat vaginal blood flow can be driven by 20-30 minutes of direct pudendal and tibial nerve stimulation [23], [24]. This stimulation effect may lead to an overall improvement in vaginal and genital health, including a modulation of the vaginal microbiome, particularly with repeated sessions.

The primary hypothesis of this study was that genital arousal FSD-treatment relevant peripheral nerve stimulation modulates the vaginal microbiome. Using a non-invasive approach, we sought to bridge a potential relationship between our clinical findings and blood flow changes observed in rodents. Although rats are a standard model for sexual function research [25], it is not clear if they are a relevant animal model for the vaginal microbiome of women. Little is known about the rat vaginal microbiota. Current knowledge about the composition of the rat vaginal microbiota is based on culture-dependent studies [26]. Though total bacterial abundance has been reported to vary in relation to the rat estrous cycle [27], [28], changes in bacterial community composition based on culture-independent analysis in relation to the estrous cycle have yet to be identified. It is important to understand the composition and dynamics of the rat vaginal microbiota for consideration in reproductive, sexual function, and microbiome studies, and for determining its use as a model for humans. Thus in addition to examining the effect of nerve stimulation on the vaginal microbiome, this exploratory study investigated the composition and cycling of the healthy rat vaginal microbiota. Although stimulation did not result in a clear shift in microbiota versus a control group, some animals experienced a relative reduction in diversity, suggesting that nerve stimulation may be a treatment option for normalizing the vaginal microbiome. We also observed that the rat vaginal microbiome is dynamic and includes bacteria that fluctuate in abundance across the estrous cycle and in relation to each other. Finally, this study indicates that rats share characteristics with human vaginal microbiota and lays the groundwork for future research.

## Materials and methods

### Animals and experimental timeline

All procedures were approved by the University of Michigan Animal Care and Use Committee in accordance with the National Institutes of Health’s guidelines for the care and use of laboratory animals. Experiments were conducted in 10 nulliparous female Sprague-Dawley rats (Charles River Breeding Labs, Wilmington, MA, USA) weighing 200 to 250 g. Two groups of 5 rats apiece were selected for this study based on group sizes in two prior rat vaginal microbiome studies that took repeated samples [27], [28]. Each rat was approximately 8 weeks old at study initiation, and was healthy without dysbiosis to our knowledge. The animals were individually housed in isolated, ventilated cages to limit cross-contamination, with controlled temperature, humidity, and a 12-h light/dark cycle. Animals were provided with laboratory chow (5L0D, LabDiet, St. Louis, MO, USA) and tap water ad libitum, with enrichment provided by a plastic tube and an EnviroPak (W.F. Fisher & Son, Branchburg, NJ, USA). The 10 animals were randomly sorted into Treatment and Control groups. The experimental timeline for each animal was divided into two parts: first a two-week baseline period and then a six-week trial period. At the end of this study, each animal was used for a terminal study for other experimental objectives and then euthanized with an overdose of sodium pentobarbital (400 mg/kg).

### Lavage sampling and estrous characterization

During the intervals described below, standard procedures were followed to obtain vaginal lavage samples [29], [30]. Careful technique was used to ensure that sampling did not induce pseudopregnancy in the animals, which is known to have an effect on the estrous cycle [29]. Clean disposable gloves were used to avoid contamination with study personnel microbiota. Additionally, care was taken to avoid contamination from perineal microbiota. After brushing the perineal region with an alcohol swab, the tip of the micropipette was carefully introduced into the vagina without coming into contact with the perineal region. Three 100 μl lavage samples were obtained in sequence and combined in a plastic vial. Approximately 50 μl of the total sample was pipetted onto a glass microscope slide and visualized under a standard light microscope with 10x magnification. Characterization of the vaginal smear was used to determine the estrous stage of each sample [30]. Estrous cycle determination was performed without reference to the animal treatment group. Proestrus was identified by the presence of clusters of round nucleated epithelial cells with granular appearances. Estrus was identified by the presence of large numbers of cornified or keratinized cells which appear to have jagged edges. Metestrus was identified as a transitional stage by the presence of cornified or rounded epithelial cells as well as leukocytes. Diestrus was identified by the presence of leukocytes as well as small numbers of epithelial cells suggesting the end of diestrus and the beginning of the next cycle. Samples whose estrous determination was clearly identifiable were used for analysis. Samples whose estrous determination was unclear were not used in analysis. Further data analysis was not performed with the lavage slide samples. The remaining 250 μl of the vaginal lavage sample was transferred into a bead plate, which was kept in a −80 °C freezer until DNA isolation.

### Baseline procedure

During the baseline period, vaginal lavage samples were taken across a 14-day period to record baseline microbiota changes and relative abundances over multiple estrous cycles. An average of 10 samples were taken for each animal during this period. Baseline procedures were the same for Treatment and Control groups. Baseline sampling was performed at the same time of day for each animal to limit the possibility of mischaracterizing the estrous phase during transition periods [31]. The animals were anesthetized with 5% Isoflurane in oxygen. Vaginal lavage samples were taken for each animal, at which point Isoflurane was turned off and the animals were allowed to wake up on their own. Samples were characterized under the microscope as described above and transferred to the beaded well plate for storage until sequencing.

### Experimental procedures

Trial sessions were performed twice a week for six weeks. Both Treatment and Control animals were anesthetized with intraperitoneal injections of ketamine and xylazine (50 mg/kg, 5 mg/kg respectively) in a cocktail diluted to 20 mg/ml and 2 mg/ml respectively. Ketamine was the preferred anesthetic, as it has been used in prior studies of anesthetized rats for evaluating sexual arousal [24], [32], [33]. Respiratory rate and skin temperature were monitored every five minutes for the duration of the procedure to ensure that the animal remained properly sedated. Respiratory rate was monitored by counting chest contractions, and skin temperature was monitored using a laser thermometer (IRT-2, Thermoworks, Salt Lake City, Utah, USA).

Each animal was placed on its back onto a surgical pad. Once visibly sedated, a toe pinch was performed to ensure that the rat had no visible response to pain sensation. A pre-trial vaginal lavage was taken, estrous phase was visualized under the microscope, and the sample was transferred to the beaded well plate. A sterilized bipolar stimulation hook (PNB0.8×2/90, Xi’an Friendship Medical Electronics, Xi’an, Shaanxi, China) was placed with the top hook directly on the clitoris and the bottom hook just below the vaginal opening. The hook was held in place by a micromanipulator (KITE-M3-R, World Precision Instruments, Sarasota, FL, USA). The probe was connected to a stimulus pulse generator (4100, AM Systems, Sequim, WA, USA) which was set to biphasic, current-controlled stimulation.

To set the stimulation amplitude, pudendo-pudendal reflex activation of the external anal sphincter was identified in Treatment animals [34]. The pulse generator was set to a pulse width of 1000 μs, a frequency of 1 Hz, and an amplitude of 2.0 mA in order to test for a visible response. The probe position was adjusted until a reflex anal sphincter response was identified in order to ensure that the probe was making proper contact with the genital nerve region. The frequency was then increased to 10 Hz and the amplitude was reduced to 0.1 mA. The amplitude was steadily increased until a muscle response was observed and then the amplitude was lowered just below that threshold. Subsequently, stimulation was applied continuously for 30 minutes. Sterile gauze was used to wipe away any urine that accumulated during the 30-minute session as to not interfere with the electrical conduction. Control animals were positioned just as Treatment animals, with the hook in place, but stimulation was not delivered. Treatment and Control animals were taken off anesthesia using an intramuscular injection of antisedan (5 mg/ml) and placed in individual cages until recovery, at which point they were transferred back to their housing cages.

### Data sequencing and microbiome analysis

DNA isolation, library preparation, and sequencing were performed by the University of Michigan Microbial Systems Molecular Biology Laboratory. DNA was isolated in an Eppendorf EpMotion 5075 liquid handling system, using a MagAttract PowerMicrobiome DNA/RNA Kit (27500-4 EP/27500-4 EP-BP, Qiagen, Hilden, Germany). Amplicon library preparation and sequencing were done as described previously [35], with the following modifications to the PCR. Briefly, the V4 region of the 16S rRNA gene was amplified from 5 or 7 µl DNA by standard PCR (described in [35]) using the Dual indexing sequencing strategy [36]. If amplification by standard PCR failed, 7 µl of DNA were amplified by touchdown PCR (1x(2 min at 95 °C), 20x(20 s at 95 °C, 15 s at annealing temperature (starts at 60°C, decreases 0.3°C/cycle), 5 min at 72°C), 20x(20 s at 95°C, 15 s at 55°C, 5 min at 72°C), 1x(10min at 72°C)), with a total of 40 amplification cycles rather than 30 cycles in the standard PCR. Sequencing was done on the Illumina MiSeq platform, using a MiSeq Reagent Kit V2 500 cycles (Illumina cat# MS102-2003), according to the manufacturer’s instructions with some modifications as described previously [35].

The 16S rRNA gene sequence data was processed and analyzed using the software package mothur (v.1.40.2) [36], [37]. After sequence processing and alignment to the SILVA reference alignment (release 128) [38], sequences were binned into operational taxonomic units (OTUs) based on 97% sequence similarity using the OptiClust method [39]. We subsampled 1678 sequences per sample and excluded samples with fewer than 1678 sequences from our analysis. The Sequence count for each OTU was determined, and Thetayc (Θyc) distances, taking into account relative abundances of both shared and non-shared OTU, were calculated between sample communities [40]. Taxonomic classifications of OTUs were based on Ribosomal Database Project (RDP) training set (version 16) [41], [42].

### Statistical analysis

Two forms of data were used in statistical analysis: relative abundance of each OTU, and Θyc distances. The sequence counts for each OTU had excessive zeros, as is common in microbiome studies [43], and causes most OTU data to not be distributed normally [44], In keeping with this assumption of non-normality, the sequence count data was analyzed using nonparametric statistical tests [45]. Samples were identified as either control or treatment, baseline or trial, and by stage of estrous cycle. Community compositions bar plots were made to visualize the relative abundance of the most abundant taxa for each animal over the baseline and trial periods. Principal coordinate analysis (PCoA) [46] was used to visualize Θyc distances between samples and to observe potential clustering of baseline samples based on estrous characterization. Analysis of Molecular Variance (AMOVA) [47] of Θyc distances was used to detect significant differences between the composition of baseline samples based on estrous characterization. Kruskal Wallis analysis was used to detect significant differences in the abundance of each OTU based on estrous characterization. The significance level α was set to 0.05, but Mann Whitney post hoc analyses were run on all samples with a *p*-value < 0.1 to also evaluate samples that showed a trend toward statistical significance. Linear Discriminant Analysis (LDA) Effect Size (LEfSe) was used to confirm differential abundance of OTU based on estrous stage [48]. Correlation analysis was run to determine if the relative abundance of pairs of OTU was significantly correlated over time.

To evaluate the effects of stimulation on sample diversity, first an AMOVA of Θyc distances between Control and Treatment animals at each time point was performed. Mann Whitney analysis was used to detect significant changes in the abundance of each OTU between baseline and trial periods. Kruskal Wallis and Mann Whitney post hoc tests were used to determine if there were changes in OTU abundance based on estrous cycling between baseline and trial periods. Lastly, Θyc distances between consecutive samples were plotted across each animal’s experimental timeline to determine the changes in diversity from one sample to the next across time. Mean Θyc distances were calculated for baseline and trial periods and a Trial:Baseline ratio was used to determine mean changes in the diversity between baseline and trial periods for each animal. Chi Squared analysis was used to detect significant variance within Control and Treatment groups. Kruskal Wallis, Mann Whitney, and Spearman tests were run using SPSS (IBM SPSS Statistics for Windows, Version 25.0. IBM Corp, Armonk, NY, USA). Community compositions bar plots, PCoA, and sequence count boxplots were made using R (Version 3.4.4. R Foundation for Statistical Computing, Vienna, Austria). AMOVA and LEfSe analyses were run using mothur (Version 1.40.2, University of Michigan, Ann Arbor, Michigan, USA). The significance level α for all tests was set at 0.05.

## Results

### Vaginal bacterial community composition of rats

Six Treatment animals and four Control animals were used in this study. One treatment animal died after baseline sampling due to complications from anesthesia and its data was only used for baseline analysis. After sequence processing and exclusion of samples with fewer than 1678 sequences, 306 samples were included in the analysis. Each of the nine full-study animals provided a mean sample count of 32.8 ± 1.45 samples (range 30-34), excluding the animal used only for baseline analysis, which only provided 10 samples. All sequence data can be found at the National Center for Biotechnology Information (NCBI) BioProject repository (Accession PRJNA545958).

A total of 1,591 different OTUs were identified in the dataset. The first twenty OTUs were focused on in analyses, as OTUs 1-14, 16, 17, 19, and 20 each had at least 10 counts in at least 10% of the samples [49]. Taxonomic nomenclature for the first twenty OTUs is reported in Table 1. The confidence of the taxonomic ranking for these first twenty OTU was 100%. The community composition of each sample, representing the bacterial phyla and genus level rankings for major OTU present in the animals in the study are shown in Fig 1. The most abundant phyla across all animals were *Proteobacteria, Firmicutes*, and *Actinobacteria*. The relative abundances of these phyla appeared to change over time for each animal during baseline and trial periods, suggesting a dynamic microbiome.

**Table 1.** Taxonomic ranking of twenty most abundant OTU.

| OTU | Kingdom | Phylum | Class | Order | Family | Genus |
| --- | --- | --- | --- | --- | --- | --- |
| OTU1 | *Bacteria* | *Proteobacteria* | *Gammaproteobacteria* | *Enterobacteriales* | *Enterobacteriaceae* | *Proteus* |
| OTU2 | *Bacteria* | *Proteobacteria* | *Gammaproteobacteria* | *Enterobacteriales* | *Enterobacteriaceae* | *Escherichia/Shigella* |
| OTU3 | *Bacteria* | *Firmicutes* | *Bacilli* | *Lactobacillales* | *Streptococcaceae* | *Streptococcus* |
| OTU4 | *Bacteria* | *Proteobacteria* | *Gammaproteobacteria* | *Enterobacteriales* | *Enterobacteriaceae* | *Morganella* |
| OTU5 | *Bacteria* | *Proteobacteria* | *Gammaproteobacteria* | *Pasteurellales* | *Pasteurellaceae* | *Pasteurellaceae* |
| OTU6 | *Bacteria* | *Firmicutes* | *Bacilli* | *Lactobacillales* | *Enterococcaceae* | *Enterococcus* |
| OTU7 | *Bacteria* | *Actinobacteria* | *Actinobacteria* | *Actinomycetales* | *Corynebacteriaceae* | *Corynebacterium* |
| OTU8 | *Bacteria* | *Firmicutes* | *Bacilli* | *Lactobacillales* | *Streptococcaceae* | *Streptococcus* |
| OTU9 | *Bacteria* | *Firmicutes* | *Bacilli* | *Lactobacillales* | *Aerococcaceae* | *Aerococcus* |
| OTU10 | *Bacteria* | *Firmicutes* | *Bacilli* | *Bacillales* | *Streptococcaceae* | *Jeotgalicoccus* |
| OTU11 | *Bacteria* | *Firmicutes* | *Bacilli* | *Bacillales* | *Streptococcaceae* | *Staphylococcus* |
| OTU12 | *Bacteria* | *Firmicutes* | *Bacilli* | *Lactobacillales* | *Aerococcaceae* | *Facklamia* |
| OTU13 | *Bacteria* | *Firmicutes* | *Clostridia* | *Clostridiales* | *Peptostreptococcaceae* | *Romboutsia* |
| OTU14 | *Bacteria* | *Firmicutes* | *Bacilli* | *Bacillales* | *Bacillaceae* | *Bacillaceae* |
| OTU15 | *Bacteria* | *Firmicutes* | *Bacilli* | *Lactobacillales* | *Lactobacillaceae* | *Lactobacillus* |
| OTU16 | *Bacteria* | *Firmicutes* | *Bacilli* | *Lactobacillales* | *Carnobacteriaceae* | *Atopostipes* |
| OTU17 | *Bacteria* | *Firmicutes* | *Erysipelotrichia* | *Erysipelotrichales* | *Erysipelotrichaceae* | *Turicibacter* |
| OTU18 | *Bacteria* | *Firmicutes* | *Negativicutes* | *Selenomonadales* | *Veillonellaceae* | *Veillonella* |
| OTU19 | *Bacteria* | *Firmicutes* | *Bacilli* | *Lactobacillales* | *Streptococcaceae* | *Streptococcus* |
| OTU20 | *Bacteria* | *Verrucomicrobia* | *Verrucomicrobiae* | *Verrucomicrobiales* | *Verrucomicrobiaceae* | *Akkermansia* |

**Fig 1.**
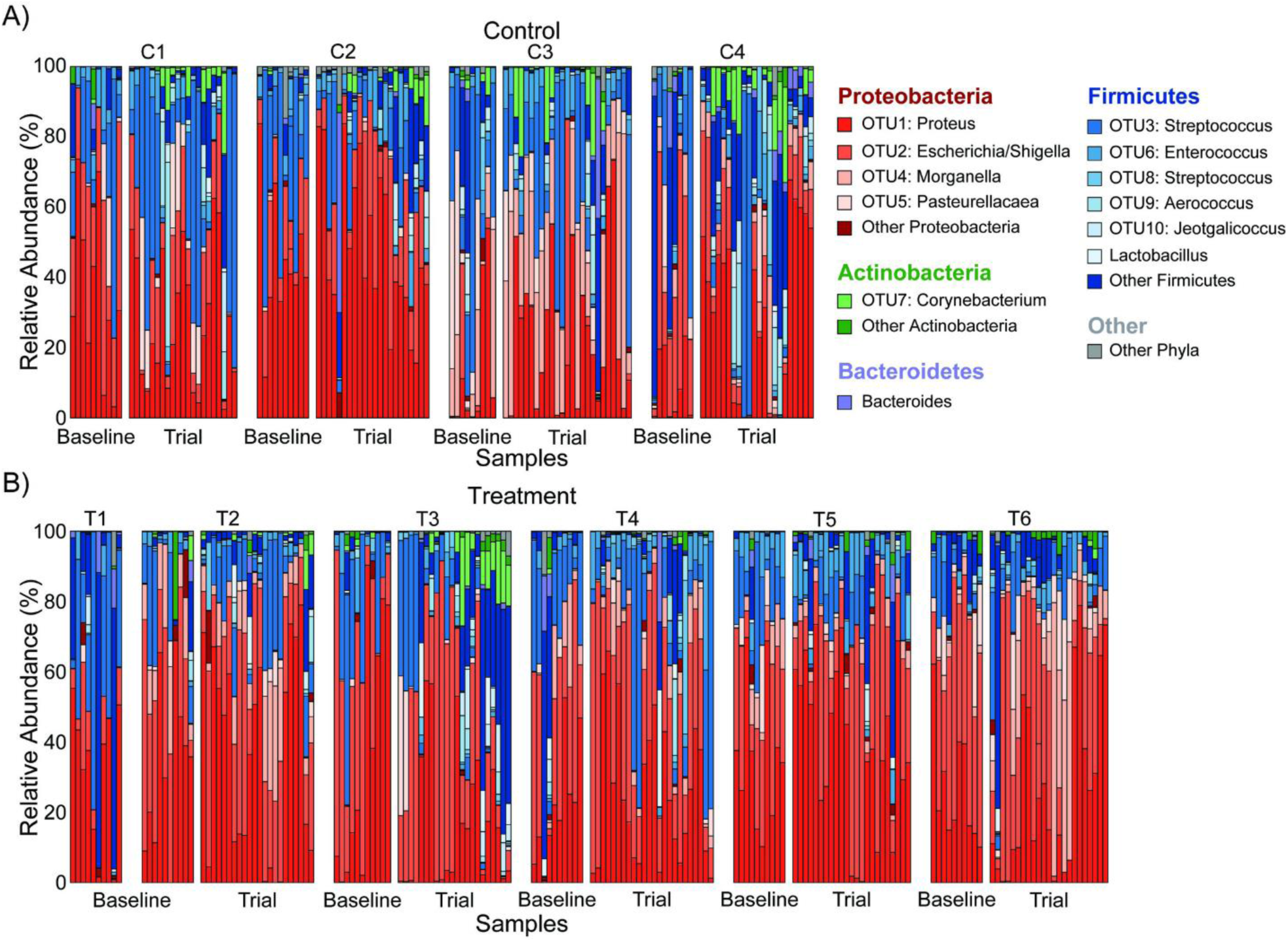
Community composition of Control and Treatment samples over time. Changes in relative abundance of key groups plotted over the baseline and trial periods for A) Control and B) Treatment animals.

### Association between estrous cycle and rat vaginal microbiota

AMOVA detected a significant difference in the Θyc distances between samples based on the estrous phase (Fs = 2.57, *p* = 0.005). Pairwise analysis showed a significant difference between estrus and diestrus samples (Fs = 5.29, *p* = 0.002) and a near-significant difference between metestrus and diestrus samples (Fs = 2.28, *p* = 0.059). Other comparisons were not significant (Fs = 1.01–2.09, *p* = 0.111–0.122). The Θyc distances between samples during the baseline period was visualized by Principal Coordinate Analysis (PCoA) (Fig 2). Biplot arrows for the ten OTUs with the lowest axis 1 *p*-values, suggesting the greatest variation between samples, are also indicated. In PCoA, samples showed a degree of clustering during estrus and diestrus, though no clear sign of clustering was apparent during proestrus and metestrus. Samples in estrus clustered around OTU3 (*Streptococcus*), while samples in diestrus clustered around OTU1 (*Proteus*) and OTU2 (*Escherichia*/*Shigella*), suggesting that OTU3 is more abundant on estrus and that OTU1 is more abundant during diestrus.

**Fig 2.**
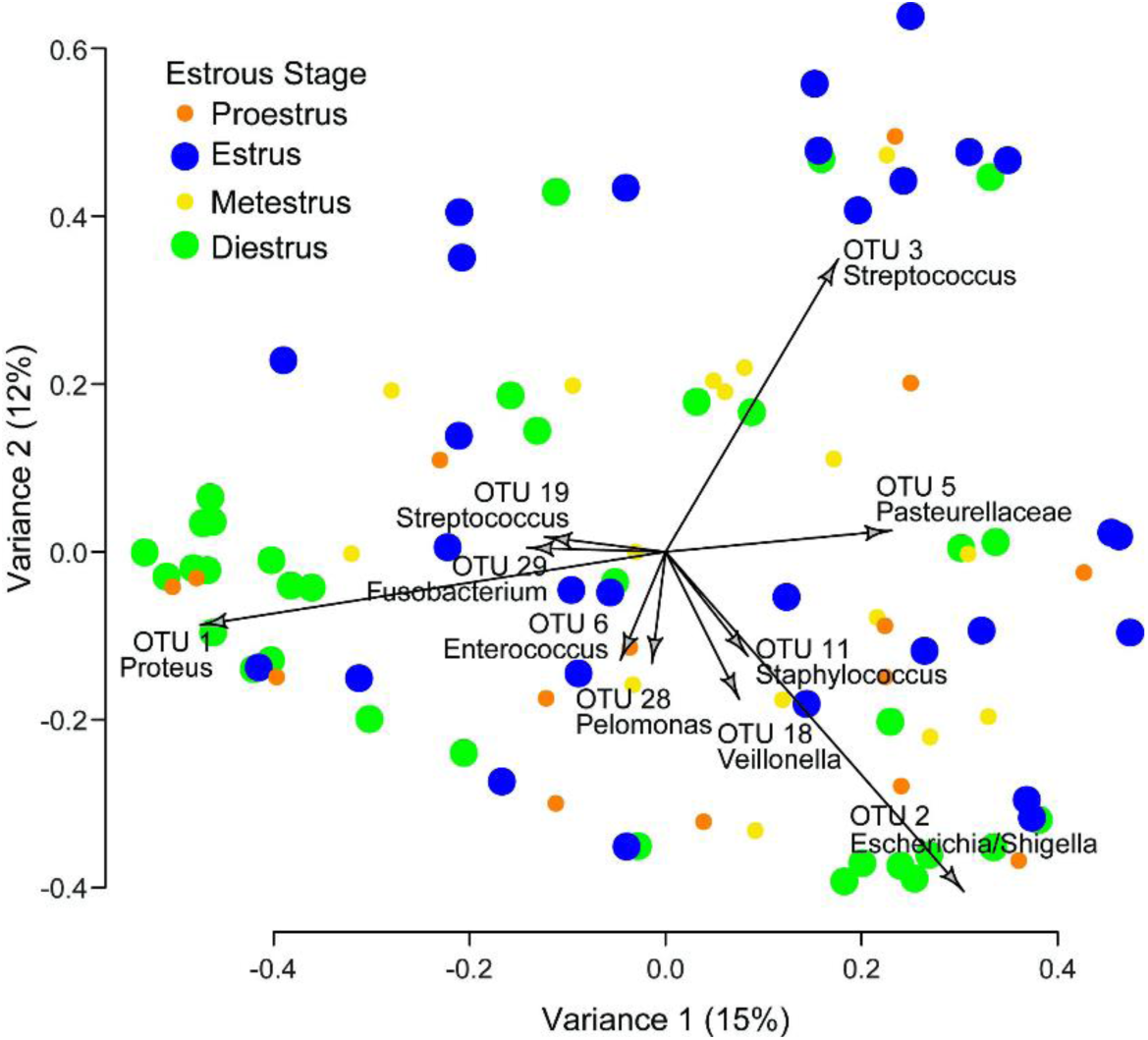
Principal Coordinate Analysis (PCoA) of all samples during baseline period. Two-dimensional distances identify dissimilarities between samples based on relative abundances of OTU. Biplot arrows indicate specific OTU which drive samples to different locations on the plot. The first PCoA axis (x-axis) included 15% of all variance. The second PCoA axis (y-axis) included 12% of all variance. Icons for estrus and diestrus are enlarged to help visualize the relative clustering of each.

After determining that the microbiota composition may change based on the estrous cycle, tests were run to determine if specific OTU had varying abundances based on the estrous stage using both Control and Treatment samples during the baseline period. Control and Treatment animals were not combined during the Trial period due to the possible effect of stimulation. Linear Discriminant Analysis Effect Size (LEfSe) tests were run and showed significant differences for only OTU3 (*Streptococcus*) and OTU7 (*Corynebacterium*) across the estrous cycle. OTU3 (*Streptococcus*) was significantly more abundant during estrus compared to any of the other three stages (LDA = 5.182, *p* = 0.0001). OTU7 (*Corynebacterium*) was significantly more abundant during diestrus compared to any of the other three stages (LDA = 4.543, *p* = 0.0003). These trends are shown in Figure 3.

**Fig 3.**
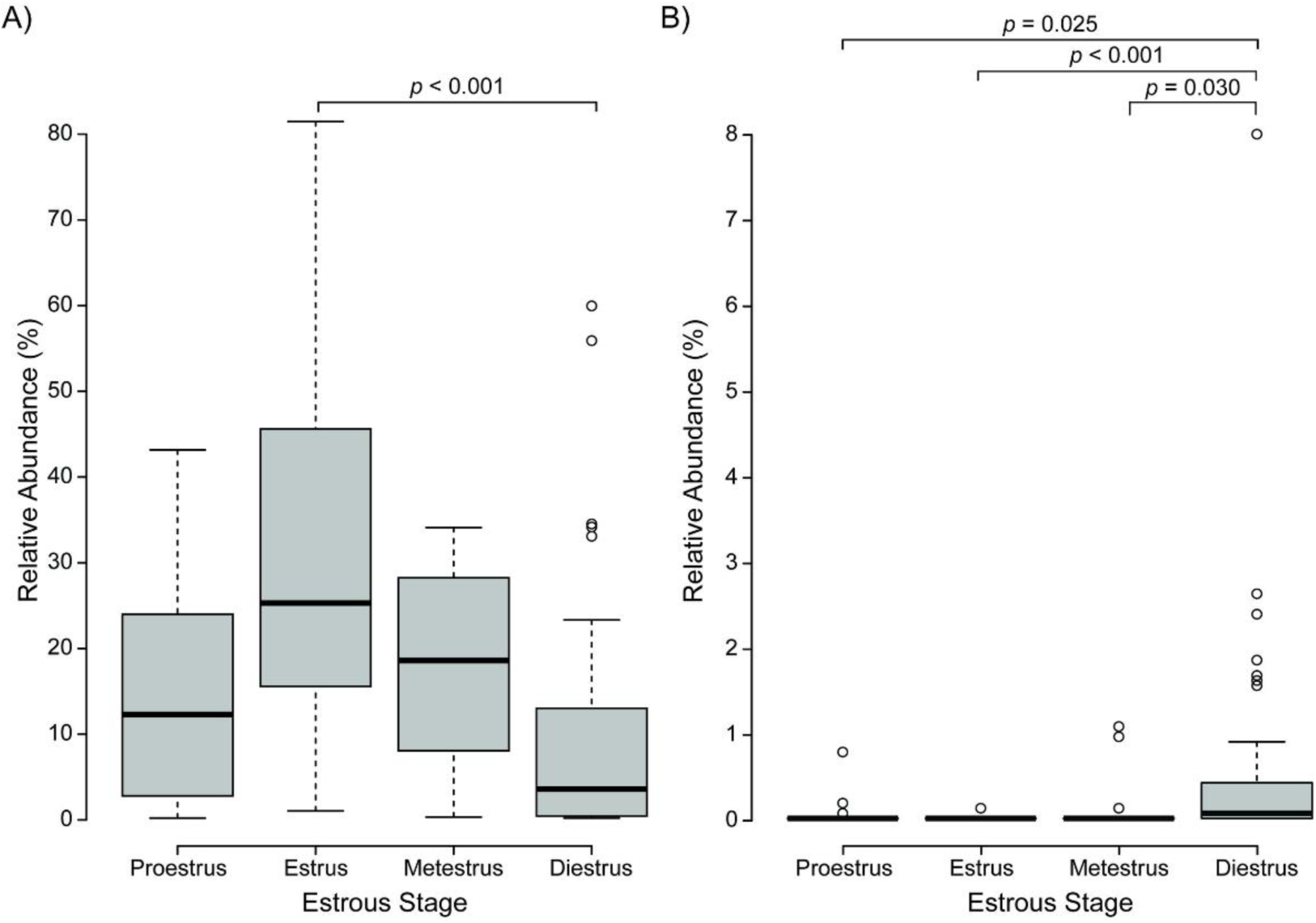
Variations in OTU abundances based on estrous phase characterization. A) OTU3 (*Streptococcus*) for Control and Treatment animals during baseline period. B) OTU7 (*Corynebacterium*) for Control and Treatment animals during baseline period. Similar trends were observed for both OTU3 and OTU7 when separated into Control-only and Treatment-only analyses.

To better understand how the microbiome can change over time, it was important to also understand how OTU abundances change in relationship to one another. For this, Spearman correlation tests were run for each combination of the first 20 OTUs over time. Correlations were determined for both Control and Treatment groups during the baseline, trial, and combined baseline & trial periods (S1 Table). Comparisons were not performed between Control and Treatment animals during the Trial period due to the possible effect of stimulation. Three example OTU-pair correlations showing typical strong relationships are given in Fig 4. Across all of the paired OTU combinations there was variability in relationship strength (R/*p*-value) and direction (positive/negative), both for pairs in the different animal-set groupings and also for individual OTUs against other OTUs. Some trends were evident. OTU3 (*Streptococcus*) and OTU6 (*Enterococcus*) more consistently had inverse relationships to other OTUs, such that their relationship to another OTU was more likely to be negative for all seven datasets in S1 Table. OTU10 (*Jeotgalicoccus*) and OTU17 (*Turicibacter*) were the most consistent in having direct relationships with other OTUs, with OTUs 7-9, 11-13, and 20 just below. OTUs 7, 9-11, 12-17, and 20 were the most likely to have the strongest relationships with other OTUs, with 26-41% of their respective combinations having *p* < 0.001.

**Fig 4.**
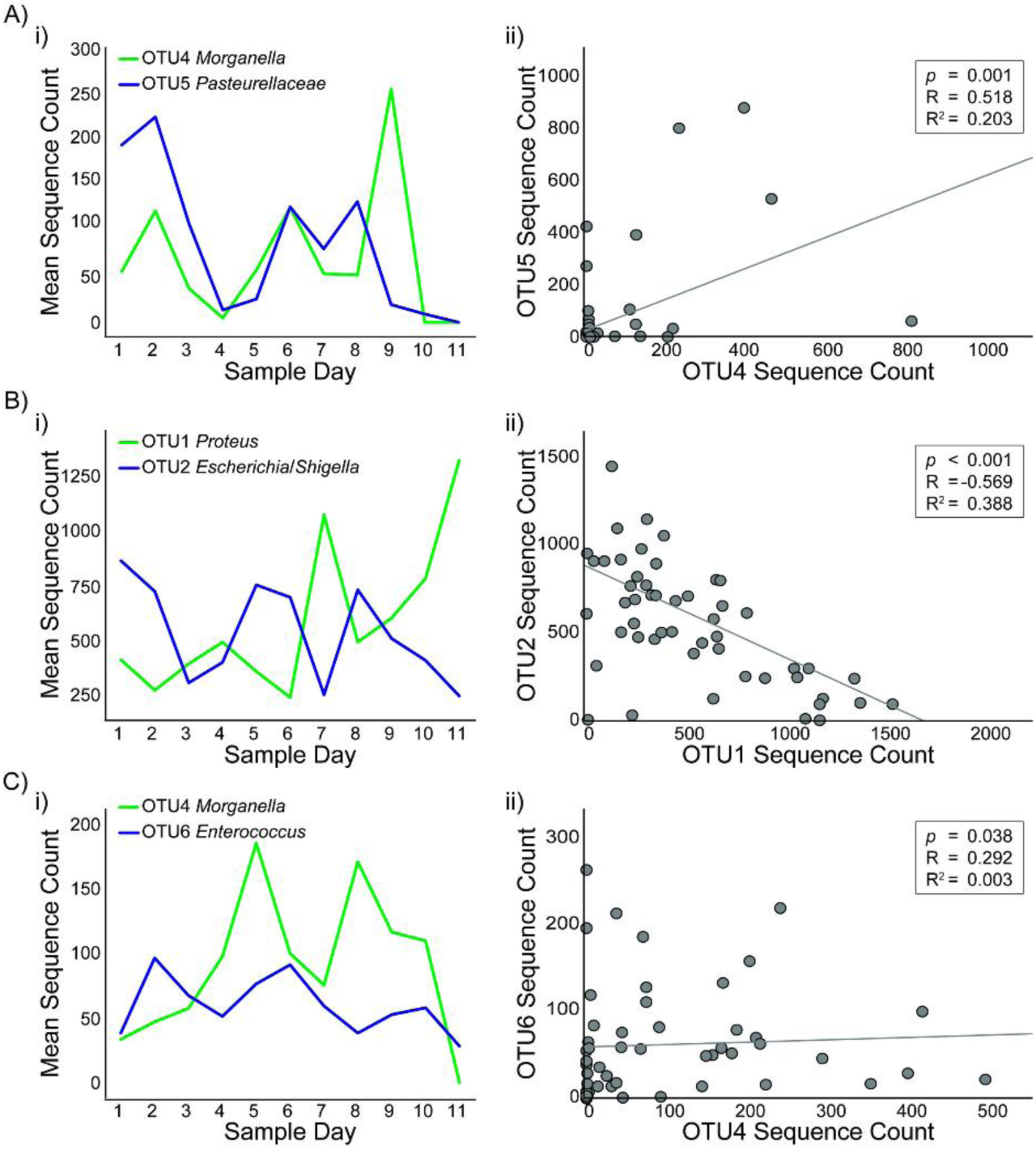
Examples of strong correlations in paired OTU sequence counts during baseline period. A) Strong negative correlation between OTU1 (*Proteus*) and OTU5 (*Pasteurellaceae*) for all animals i) mean relative abundances and ii) paired scatter plot. B) Strong positive correlation between OTU9 (*Aerococcus*) and OTU10 (*Jeotgalicoccus*) in all animals, i) mean relative abundances and ii) paired scatter plot. C) Strong positive correlation between OTU13 (*Romboutsia*) and OTU17 (*Turicibacter*) in all animals, i) mean relative abundances and ii) paired scatter plot.

### Stimulation effects

To determine the potential effects of stimulation on the microbiome diversity, three tests were run. An AMOVA analysis of Θyc distances between Control and Treatment animals at each study time point indicated that two time points during baseline (Baseline 7: Fs = 3.02, *p* = 0.036; Baseline 8: Fs = 5.83, *p* = 0.01) and one date in the test period (Day 4 session: Fs = 2.65, *p* = 0.014) were different between Control and Treatment animals. There was no clear indicator as to the source of these individual differences.

Control and Treatment samples were tested for significant changes in OTU relative abundances from baseline to trial periods. Mann Whitney tests were used to evaluate changes for each animal between periods and between each phase of the estrous cycle for each of the first 7 OTU. The ratios of the Mann Whitney mean rank values for the baseline period to the mean rank values for the trial period were calculated (B:T Ratio), to determine which experimental segment showed the greatest abundance of each OTU (S2-S3 Tables). Individual OTU had significant changes in abundance from baseline to trial periods among both Control and Treatment animals and during several different days of the estrous cycle. However there was no consistent trend in overall abundance for any OTU in either Control or Treatment groups.

To determine if stimulation had longitudinal effects on the composition of the microbiota over time, Θyc distances were tracked between consecutive samples across the baseline and trial periods for Control and Treatment animals. Each sample was identified by its estrous stage characterization, with samples transitioning between estrous stage identified by an intermediate estrous stage characterization. Individual animal Θyc scatterplots did not show trends in either Control or Treatment groups, based on differences between baseline and trial periods or based on differences due to estrous stage (Fig 5A). Average Θyc distances were determined for each animal’s baseline and trial periods. To visualize the sample-to-sample variability differences between baseline and trial periods, the ratio (RT:B) of the trial average Θyc distance to the baseline average Θyc distance was then calculated (Fig 5B). The mean R_T:B_ for Control animals (1.264) and Treatment animals (1.308) was not significantly different (*p* = 0.730). All Control animals had a similar increase in mean Θyc distances from baseline to trial periods with a coefficient of variance of 0.91%. Four of the five Treatment animals had increases in mean Θyc distances, though the degree of change was more variable than Control animals, with a coefficient of variance of 29.4%. The variance in R_T:B_ among Treatment animals was significantly higher than that of Control animals, as confirmed by a chi squared analysis (*p* = 0.00005).

**Figure 5.**
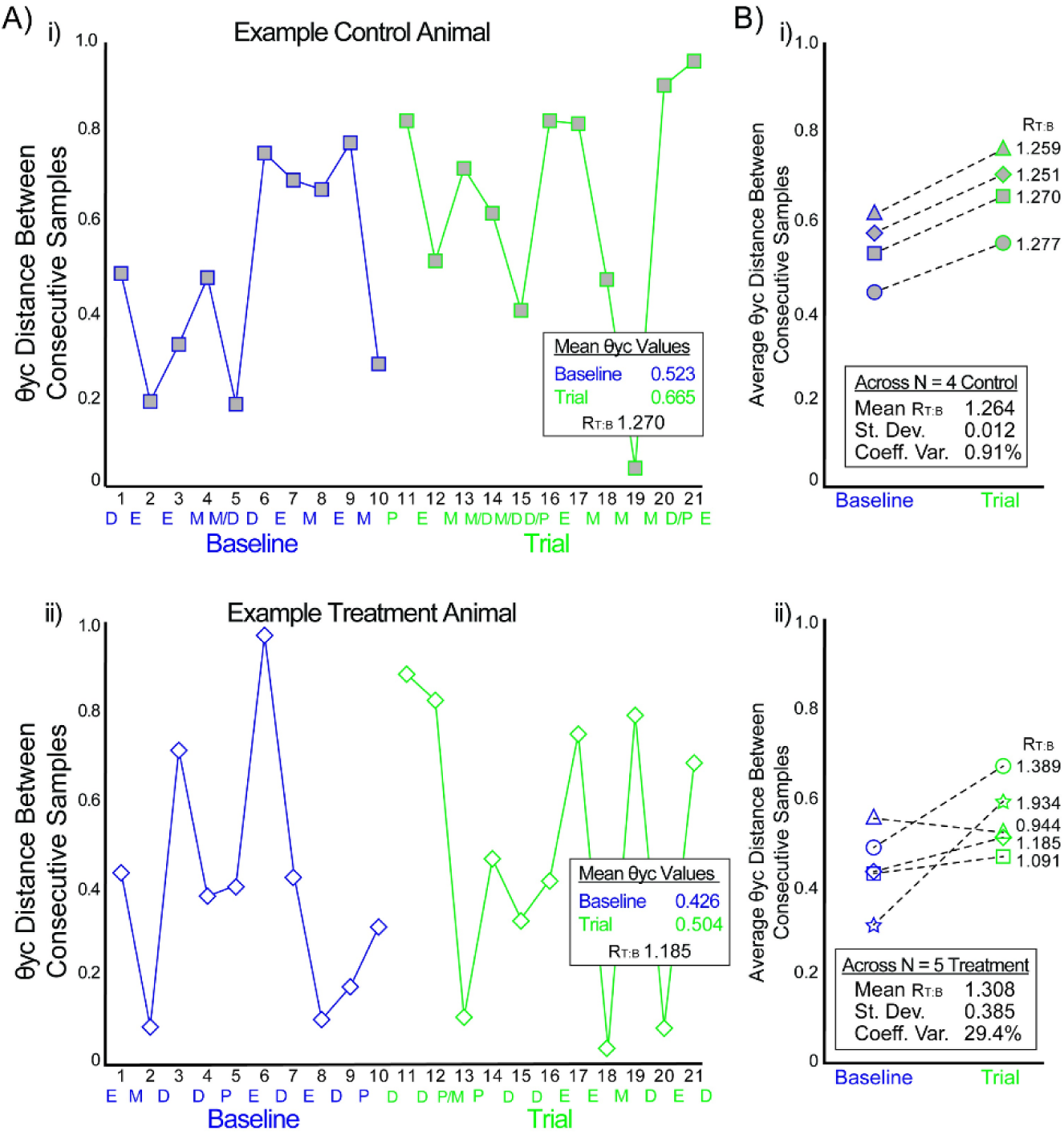
Θyc distances between consecutive samples for Control and Treatment groups. A) Example scatterplots of Θyc distances for consecutive samples for i) Control 1and ii) Treatment 2 animals. Baseline period in blue; Trial period in green. Estrous period (P/E/M/D for states) indicated at bottom of each plot. B) Mean Θyc distances during baseline and trial periods for i) Control and ii) Treatment animals. Icons in (A) correspond to same animals in (B).

## Discussion

To our knowledge, this is the first study to explore the complexity of the changing vaginal microbiota composition in rats across the estrous cycle. Previous studies have made broad claims about the overall abundance of bacteria during the estrous cycle [50], but this study is the first to identify changes in specific OTU relative abundances and identify correlations between OTUs across time. Our results suggest a dynamic vaginal microbiome (Table 1, Fig 1) that shifts based on both changes in specific OTU relative abundances by estrous stage (Fig 2-3) and in correlations among the OTUs (Fig 4, S1 Table). Additionally, this is the first study to our knowledge that examines the effect of neural stimulation on the microbiome of any organ in any species. Specifically, this study evaluated the effect of a specific form of neural stimulation on the vaginal microbiome of healthy rats. Though the effect of stimulation was ambiguous (S2-S3 Tables), analysis of Θyc changes over time showed differences in variance among Treatment animals when compared to Control animals (Fig 5), suggesting a potential effect of stimulation on vaginal microbiome diversity.

Several statistical tests confirmed that specific OTU relative abundance levels varied based on the stage of the estrous cycle. Initial community compositions showed that OTU relative abundance can change across relatively short time periods (Fig 1). PCoA clustering suggested that the changing of community composition may be related to the estrous cycle (Fig 2). Certain OTUs were more abundant during specific estrous stages compared to other stages. Though the total variance reported by PCoA was low (27%), current literature suggests that, even when yielding a low percent variation, clustering can yield biological insight and patterns can be revealed [46]. This was confirmed by AMOVA analysis of Θyc which showed that the Θyc distances, which correlate to the overall difference in community composition between samples, were significantly different between estrus and diestrus. Clearly, estrous cycling in the rat vagina has an effect on the overall composition of the microbiome.

Further analysis was done to identify which specific OTUs showed significant differences in relative abundance based on the estrous cycling. The LEfSe analysis showed that OTU3 (*Streptococcus*) was significantly more abundant during estrus than any other stage and OTU7 (*Corynebacterium*) was significantly more abundant during diestrus than any other stage. Together, this suggests that a factor in the overall difference in bacterial abundance between estrous stages [27] is not a global increasing or decreasing of all bacterial abundances but may be due to specific OTU adjusting in relative abundance based on the estrous cycle.

Changes in the vaginal microbiome due to estrous cycling are thought to be correlated to several factors such as pH, hormones, presence of mucin, and other biological conditions [27], [51]. During estrus, mucus secretion from glands located in the cervix of the uterus increase the mucus in the environment [52], [53]. It is theorized that the microbiome flourishes during this time because the mucus acts as a medium for bacterial growth [27]. Estrus would appear to be the optimal estrous stage for bacterial growth, suggesting that there is a global increase in bacterial composition during estrus. This assumption, however, does not take into account the dynamics of each OTU. Our study suggests that while some species favor estrus, other favor diestrus. Several OTUs, for example, showed predominance during diestrus compared to estrus. OTU1 (*Proteus*) showed clustering in PCoA during diestrus (Fig 2), and OTU7 (*Corynebacterium*) was significantly more abundant during diestrus compared to estrus across multiple forms of analysis (Fig 3). This would suggest that several OTU, including OTU1 (*Proteus*), the most abundant OTU found in our study, tend to favor diestrus. It was also interesting to observe that the vast majority of differences in OTU abundance occurred between estrus and diestrus, suggesting that proestrus and metestrus act as transitions between points of significant bacterial growth or decay. While the data do not show that all of the OTU were significantly different in abundance across the estrous cycle, it is theorized that most of the OTU are changing slightly from estrous stage to stage and certain OTUs change more dramatically than others. It is not understood why specific OTUs showed significant differential abundances depending on the estrous state and other OTUs did not. Further studies are necessary to understand this phenomenon.

There were many significant correlations between OTUs (Fig 4, S1 Table), suggesting that OTUs can be related to each other. It was interesting to note that positive and negative correlations were found between both OTUs of similar and different taxonomic ranking, within and across the different comparison sets. For example, OTU1 (*Proteus*), OTU2 (*Escherichia*/*Shigella*), and OTU4 (*Morganella*) were all in the *Enterobacteriaceae* family. In Control animals during the trial period, a strong positive correlation was found between OTU1 (*Proteus*) and OTU2 (*Escherichia*/*Shigella*) but a strong negative correlation was found between OTU2 (*Escherichia*/*Shigella*) and OTU4 (*Morganella*). Inversely, in Control animals during the trial period, a strong positive correlation was found between OTU3 (*Streptococcus* in *Firmicutes* family) and OTU5 (*Pasteurellaceae* in *Proteobacteria* family) and a strong negative correlation was found between OTU1 (*Proteus* in *Proteobacteria* family) and OTU3. Among the OTUs with consistently positive relationships with other OTUs (e.g. OTUs 7, 9, 10, 12-17, 20), most were *Firmicutes*, with OTU7 (*Corynebacterium*) an *Actinobacteria* and OTU20 (*Akkermansia*) a *Verrucomicrobia*, that had low overall sequence counts that often tracked each other. Ultimately, no clear trend was identified suggesting that more similar OTU or more different OTU have any predetermination for positive or negative correlations (S1 Table). Further study is necessary to understand the relationship between OTU and competition for resources within the vaginal microbiome of rats.

Our study demonstrates that the rat vaginal microbiome has a dynamic microbiome which can change based on the estrous cycle. This is somewhat unique compared to other animals. Baboons, pigs, and mice have all been shown to have no difference in vaginal microbiome diversity based on ovulatory cycling [35], [54], [55]. Mice have also been shown to have no shifting of intestinal microbiome diversity due to hormone shifting during estrus [56]. Dogs, however, have been shown to have a changing vaginal microbiome based on the estrous cycle, including increased diversity among OTU during estrus [57]. In humans, small fluctuations in vaginal microbiota composition are generally reported [58], [59], though larger changes in some microbiota communities in association with the menstrual cycle have been observed [60]. Interestingly, humans do have significant differences in microbiome diversity throughout the three trimesters of pregnancy and when comparing their pre- and postpartum microbiome [61].

The bacterial composition of the rat microbiome is similar to that of the mouse [50], dog [57], and pig [54], which are dominated by *Proteobacteria, Firmicutes*, and *Actinobacteria* (Fig 1, Table 1). We observed that the rat vaginal microbiota is relatively different from the baboon vaginal microbiota, which is composed primarily *Bacteroidetes, Fusobacteria*, and *Firmicutes* [55]. When compared to the human vaginal microbiota, however, the rat microbiome had some similarities. The human vaginal microbiota is heavily predominated by *Lactobacillus* making it a unique microbiome, dissimilar to all other animal models tested thus far [62]. The human microbiota, however, can have high levels of *Corynebacterium, Escherichia*/*Shigella, Streptococcus*, and *Staphylococcus* species, as well as several others [63]. This study in rats showed a relative predominance of *Proteus*, a species that is found somewhat commonly in the human vaginal microbiome [64]. Our study did not show a high predominance of *Lactobacillus* (average relative abundance of 0.57%), though it (OTU15, OTU39, and eight additional OTUs from OTU315 and beyond) and the previously listed common human vaginal bacterial genera were found (Fig 1, Table 1). *Lactobacillus* is not uncommonly found in rat microbiome studies [26], though it is not as highly abundant as in human studies. With further research, the rat may be considered a useful model for the human vaginal microbiome. The vaginal microbiota of the rats in this study were highly concordant with a prior study using cultivation-based identification [26], which showed *Proteus, Pasteurella*, and *Lactobacillus* among others, demonstrating that culture-independent methods used here align with prior methodology.

While our study did not clearly indicate that stimulation of the genital branch of the pudendal nerve drives changes in the vaginal microbiome of healthy rats, there was some evidence suggesting that stimulation may have an effect on microbiome stability. Tests run using OTU relative abundance data showed significant changes from baseline to trial in both Control and Treatment animals (S2-S3 Tables). This suggests that OTU relative abundance changes independent of stimulation and therefore stimulation could not be concluded as having a significant effect on the abundance of specific OTU across time. This may be because OTU relative abundance by nature is a somewhat variable metric [45]. Θyc distances were used to identify more broad changes in overall composition from baseline to trial periods. Again, both Control and Treatment groups showed similar changes in Θyc distances from baseline to trial periods. The mean value of change in Θyc distances from baseline to trial period did not differ significantly when comparing Control and Treatment groups (Fig 5). What was noteworthy, however, was that the coefficient of variance for Treatment animals was 32.3 times larger than for the Control animals, representing a significant difference between the groups. This suggests that stimulation may have affected the overall diversity, though without a clear trend in our individual samples or OTU.

Interestingly, while all of the Control animals and 4 of the 5 Treatment animals had an increase in the mean Θyc during the trial period, one Treatment animal had a decrease and the increase in two rats was at a lower rate than Control animals (Fig 5B). In humans, a less diverse vaginal microbiome is considered more stable and healthier [65]. The human microbiome is said to be healthy when predominated by one or a few bacterial species, such as *Lactobacillus* [66]. In a similar vein, studies have shown that women with bladder issues such as urge urinary incontinence have more diverse bladder microbiomes than women without incontinence [67], [68]. Additionally, studies have shown that women with imbalances in *Lactobacillus* in their vaginal microbiome can have changes in hormones associated with cervicovaginal inflammation, which leads to changes in their vaginal immunity and HIV susceptibility [10], [58]. It is possible that the handling and stress of the study contributed to an increase in the microbiome diversity for Control animals, and that the three Treatment rats with lower rate of increase or a decrease were affected by stimulation. Partial population responses to neuromodulation are often reported in the literature [69] and so a larger study size or use of an animal model of VVA may be useful to examine these relationships further. If genital nerve stimulation can modulate vaginal microbiome diversity, it offers hope as a treatment for VVA and suggests underlying mechanisms for its potential use in treating genital arousal aspects of female sexual dysfunction.

A possible limitation of this study was the use of excessive zeros in the data analysis. Excessive zeros in microbiome studies are common and can potentially skew data [43]. Zeros come in multiple forms [70]. Outlier zeros are due to extraneous conditions, structural zeros are due to the nature of different experimental groups, and sampling zeros are any other zero that may be due to low sampling depth. Each type of zero can be addressed differently by removing them, using pseudo counts, and other algorithms to minimize data skew [71]. This study used nonparametric statistical tests to control for non-normal distributions but did not transform or remove any of the zeros in the data set. Further studies in this field may attempt to perform analysis of microbiome data while controlling for all three types of zeros to determine if this affects the results.

While frozen storage of samples does not have an effect on microbiome diversity [72], thawing and refreezing can have a significant effect [73]. Therefore, it was important to prevent this occurrence during the experimental periods prior to sequencing. While appropriate steps were taken, it is possible that some samples may have experienced temperature changes which may have affected their bacterial composition. A maternal effect can create changes in the microbiome due to parenting techniques, which can last over several generations [46]. Though changes in individual animal microbiome were not specifically controlled for maternal effect in this study, we used young rats that had not spent time with their mothers post-weaning. Therefore, while multi-generation effects may have had a role, a first-generation maternal effect was not a limitation in this study.

This study used transcutaneous electrical stimulation, which only involves skin-contact stimulation. Prior to each stimulation interval we verified that nerve recruitment was occurring, and this skin-surface approach did not require the stress of surgery for an implant. However, our prior rat studies that observed increases in vaginal blood flow used cuff electrodes implanted directly on the pudendal [23] and tibial nerves [24]. Furthermore, while some nerve stimulation studies for bladder function have used skin-surface stimulation, most have shown greater efficacy using percutaneous or implanted stimulation approaches [74]. Thus our reduced-contact stimulation technique may have limited the efficacy of the nerve recruitment and contributed to the relatively inconclusive stimulation results in Treatment animals. Additionally, our recent clinical study using skin-surface stimulation had greater improvements in sexual function for tibial nerve stimulation than genital nerve stimulation, in a small sample size [22], suggesting that alternate nerve targets may have stronger effects. Future studies with direct neural stimulation comparing genital or pudendal nerve stimulation to other nerve targets may yield more conclusive results.

This study used healthy, nulliparous rats. We presume that their vaginal microbiota started at a healthy state, and as such the overall effects of nerve stimulation may have been limited. Ovariectomized rats are a standard animal model for postmenopausal changes including vaginal atrophy [75]. Prior research with ovariectomized rats has shown that hormone replacement treatments can ameliorate the induced vaginal atrophy [76], [77]. Future studies using nerve stimulation and hormone replacement in ovariectomized rats may be insightful into the effects of stimulation on an abnormal vaginal microbiome and allow for comparison between these treatment approaches.

## Conclusions

This study demonstrates that the rat vaginal microbiota can change based on the estrous cycle. The microbiome is dynamic, changes in diversity over time, and is composed of OTUs which fluctuate both in synchrony to and in opposition to one another. These findings support continued research into the rat vaginal microbiome and its relationship to the vaginal microbiome of other species including humans. While the rat vaginal microbiome was not completely similar to the human vaginal microbiome, many of the same major bacteria types were present, suggesting that rats may be a relevant animal model for humans in this area. Neural stimulation was not found to have a clear effect on the microbiome in these healthy rats, though stimulation may have modulated the microbiome diversity in a subset of animals. This study suggests the potential for further research into the use of neural stimulation to drive changes in the vaginal microbiome in states of dysbiosis and to increase vaginal function and health. Future studies may also examine additional physiological parameters in addition to the microbiome, such as vaginal pH and blood flow as well as hormones and other systemic factors, to obtain a more comprehensive perspective on their interrelationships.

## Acknowledgments

The authors thank Vincent Young, Michael Dority, Clarisse Betancourt Roman, Robert Hein, and the University of Michigan Host Microbiome Initiative for assistance with experimental planning and microbiome analysis, Lauren Zimmerman, Ahmad Jiman, Indie Rice, Nikolas Barrera, and other members of the Peripheral Neural Engineering & Urodynamics Lab for help with experiments and data collection, and Joseph Naiman for guidance on statistical planning and data analysis. This research was also supported in part through computational resources and services provided by Advanced Research Computing at the University of Michigan, Ann Arbor,

## Supporting information

**S1 Table. Correlations between sequence counts for Control and Treatment groups during the baseline, trial, and combined baseline & trial periods among the first 20 OTU.**

**S2 Table. Change in OTU relative abundance from baseline to treatment periods for Control animals.**

**S3 Table. Change in OTU relative abundance from baseline to treatment periods for Treatment animals.**

