## Supplemental Tables 1-3 for "The rodent vaginal microbiome across the estrous cycle and the effect of genital nerve electrical stimulation"

**S1 Table. Correlations between sequence counts for Control and Treatment groups during the baseline, trial, and combined baseline & trial periods among the first 20 OTU.**

| | All Samples - Baseline only | | Control – Baseline only | | Control – Trial only | | Control – Baseline & Trial | | Treatment – Baseline only | | Treatment – Trial only | | Treatment – Baseline & Trial | |
|---|---|---|---|---|---|---|---|---|---|---|---|---|---|---|
| OTU set | R | *p*-value | R | *p*-value | R | *p*-value | R | *p*-value | R | *p*-value | R | *p*-value | R | *p*-value |
| 1 & 2 | -0.221 | 0.029 | 0.083 | 0.622 | 0.452 | <0.001 | 0.308 | <0.001 | -0.433 | 0.001 | -0.336 | <0.001 | -0.364 | <0.001 |
| 1 & 3 | -0.259 | 0.010 | -0.312 | 0.057 | -0.352 | 0.001 | -0.356 | <0.001 | -0.221 | 0.090 | -0.404 | <0.001 | -0.335 | <0.001 |
| 1 & 4 | -0.190 | 0.061 | -0.516 | 0.001 | -0.169 | 0.112 | -0.239 | 0.007 | 0.036 | 0.785 | 0.189 | 0.040 | 0.135 | 0.072 |
| 1 & 5 | -0.378 | <0.001 | -0.571 | <0.001 | -0.296 | 0.005 | -0.379 | <0.001 | -0.225 | 0.083 | -0.355 | <0.001 | -0.306 | <0.001 |
| 1 & 6 | 0.170 | 0.094 | 0.461 | 0.004 | 0.358 | 0.001 | 0.378 | <0.001 | 0.035 | 0.792 | 0.302 | 0.001 | 0.209 | 0.005 |
| 1 & 7 | 0.061 | 0.549 | 0.148 | 0.375 | -0.110 | 0.301 | -0.004 | 0.961 | 0.052 | 0.695 | 0.031 | 0.742 | 0.025 | 0.744 |
| 1 & 8 | -0.075 | 0.461 | -0.127 | 0.446 | -0.101 | 0.345 | -0.083 | 0.354 | -0.039 | 0.765 | -0.158 | 0.088 | -0.110 | 0.143 |
| 1 & 9 | 0.084 | 0.411 | 0.147 | 0.379 | -0.088 | 0.408 | 0.014 | 0.872 | 0.060 | 0.651 | -0.034 | 0.715 | -0.025 | 0.736 |
| 1 & 10 | 0.072 | 0.482 | 0.182 | 0.274 | -0.059 | 0.581 | 0.026 | 0.768 | 0.031 | 0.815 | 0.063 | 0.497 | 0.042 | 0.581 |
| 1 & 11 | -0.095 | 0.352 | 0.060 | 0.721 | 0.236 | 0.025 | 0.186 | 0.035 | -0.213 | 0.102 | 0.064 | 0.492 | -0.006 | 0.933 |
| 1 & 12 | -0.038 | 0.709 | 0.022 | 0.897 | -0.092 | 0.390 | -0.031 | 0.727 | -0.049 | 0.709 | 0.022 | 0.810 | 0.003 | 0.966 |
| 1 & 13 | -0.090 | 0.378 | -0.172 | 0.302 | 0.068 | 0.526 | 0.011 | 0.906 | -0.021 | 0.874 | 0.017 | 0.853 | -0.001 | 0.991 |
| 1 & 14 | 0.019 | 0.854 | 0.059 | 0.724 | -0.165 | 0.121 | -0.099 | 0.268 | 0.079 | 0.549 | 0.034 | 0.713 | 0.028 | 0.706 |
| 1 & 15 | -0.263 | 0.009 | -0.383 | 0.018 | 0.048 | 0.656 | -0.046 | 0.608 | -0.123 | 0.348 | -0.080 | 0.391 | -0.107 | 0.156 |
| 1 & 16 | 0.050 | 0.627 | 0.043 | 0.798 | -0.187 | 0.077 | -0.112 | 0.208 | 0.162 | 0.217 | 0.114 | 0.219 | 0.107 | 0.156 |
| 1 & 17 | -0.139 | 0.172 | -0.261 | 0.114 | 0.013 | 0.902 | -0.050 | 0.574 | -0.049 | 0.710 | -0.036 | 0.697 | -0.051 | 0.496 |
| 1 & 18 | -0.052 | 0.609 | -0.232 | 0.160 | * | * | -0.131 | 0.141 | -0.052 | 0.693 | 0.002 | 0.982 | -0.009 | 0.903 |
| 1 & 19 | 0.126 | 0.216 | -0.114 | 0.494 | -0.013 | 0.906 | -0.012 | 0.889 | 0.290 | 0.025 | 0.187 | 0.043 | 0.210 | 0.005 |
| 1 & 20 | -0.186 | 0.067 | -0.332 | 0.042 | -0.010 | 0.924 | -0.107 | 0.231 | -0.047 | 0.722 | -0.073 | 0.434 | -0.076 | 0.316 |
| 2 & 3 | -0.113 | 0.269 | -0.341 | 0.036 | -0.237 | 0.024 | -0.178 | 0.045 | 0.026 | 0.847 | 0.112 | 0.226 | 0.094 | 0.212 |
| 2 & 4 | 0.085 | 0.408 | -0.237 | 0.153 | -0.523 | <0.001 | -0.469 | <0.001 | 0.173 | 0.187 | -0.085 | 0.358 | 0.008 | 0.911 |
| 2 & 5 | 0.145 | 0.155 | -0.186 | 0.264 | -0.239 | 0.023 | -0.191 | 0.031 | 0.324 | 0.012 | 0.137 | 0.138 | 0.214 | 0.004 |
| 2 & 6 | 0.160 | 0.115 | 0.043 | 0.800 | 0.283 | 0.007 | 0.231 | 0.009 | 0.286 | 0.027 | 0.099 | 0.288 | 0.165 | 0.028 |
| 2 & 7 | -0.141 | 0.166 | -0.109 | 0.513 | -0.104 | 0.330 | -0.209 | 0.018 | -0.091 | 0.490 | -0.144 | 0.119 | -0.135 | 0.072 |
| 2 & 8 | 0.157 | 0.122 | 0.135 | 0.419 | 0.130 | 0.221 | 0.041 | 0.648 | 0.189 | 0.149 | 0.031 | 0.739 | 0.047 | 0.537 |
| 2 & 9 | -0.123 | 0.227 | -0.137 | 0.411 | -0.059 | 0.583 | -0.185 | 0.036 | -0.063 | 0.634 | -0.189 | 0.041 | -0.168 | 0.025 |
| 2 & 10 | -0.099 | 0.331 | -0.054 | 0.749 | 0.023 | 0.833 | -0.083 | 0.354 | -0.062 | 0.637 | -0.270 | 0.003 | -0.200 | 0.007 |
| 2 & 11 | 0.234 | 0.020 | -0.033 | 0.842 | 0.302 | 0.004 | 0.180 | 0.041 | 0.311 | 0.016 | 0.010 | 0.914 | 0.123 | 0.103 |
| 2 & 12 | -0.142 | 0.162 | -0.051 | 0.759 | -0.033 | 0.756 | -0.100 | 0.260 | -0.106 | 0.418 | -0.179 | 0.053 | -0.160 | 0.033 |
| 2 & 13 | -0.316 | 0.002 | -0.033 | 0.843 | 0.190 | 0.073 | 0.078 | 0.383 | -0.443 | <0.001 | -0.248 | 0.007 | -0.327 | <0.001 |

| | All Samples - Baseline only | | Control – Baseline only | | Control – Trial only | | Control – Baseline & Trial | | Treatment – Baseline only | | Treatment – Trial only | | Treatment – Baseline & Trial | |
|---|---|---|---|---|---|---|---|---|---|---|---|---|---|---|
| OTU set | R | *p*-value | R | *p*-value | R | *p*-value | R | *p*-value | R | *p*-value | R | *p*-value | R | *p*-value |
| 2 & 14 | -0.042 | 0.678 | -0.022 | 0.896 | -0.119 | 0.265 | -0.152 | 0.086 | -0.034 | 0.797 | -0.180 | 0.051 | -0.150 | 0.046 |
| 2 & 15 | -0.124 | 0.223 | 0.026 | 0.876 | -0.070 | 0.512 | -0.092 | 0.304 | -0.206 | 0.114 | -0.287 | 0.002 | -0.260 | <0.001 |
| 2 & 16 | -0.137 | 0.179 | -0.124 | 0.459 | -0.087 | 0.413 | -0.150 | 0.091 | -0.079 | 0.549 | -0.249 | 0.006 | -0.201 | 0.007 |
| 2 & 17 | -0.305 | 0.002 | -0.042 | 0.802 | 0.221 | 0.037 | 0.100 | 0.260 | -0.409 | 0.001 | -0.303 | 0.001 | -0.337 | <0.001 |
| 2 & 18 | 0.270 | 0.007 | -0.172 | 0.301 | * | * | -0.055 | 0.535 | 0.337 | 0.008 | 0.160 | 0.083 | 0.222 | 0.003 |
| 2 & 19 | -0.193 | 0.057 | -0.343 | 0.035 | -0.320 | 0.002 | -0.347 | <0.001 | -0.114 | 0.387 | -0.076 | 0.413 | -0.103 | 0.169 |
| 2 & 20 | -0.242 | 0.016 | 0.017 | 0.921 | -0.010 | 0.926 | -0.002 | 0.979 | -0.355 | 0.005 | -0.273 | 0.003 | -0.305 | <0.001 |
| 3 & 4 | -0.035 | 0.735 | 0.087 | 0.603 | 0.028 | 0.790 | -0.001 | 0.987 | -0.079 | 0.547 | 0.147 | 0.112 | 0.041 | 0.587 |
| 3 & 5 | 0.216 | 0.033 | 0.272 | 0.098 | 0.465 | <0.001 | 0.401 | <0.001 | 0.176 | 0.179 | 0.293 | 0.001 | 0.260 | <0.001 |
| 3 & 6 | -0.048 | 0.640 | -0.269 | 0.102 | -0.225 | 0.033 | -0.222 | 0.012 | -0.013 | 0.923 | -0.092 | 0.324 | -0.068 | 0.368 |
| 3 & 7 | -0.177 | 0.081 | -0.321 | 0.049 | -0.293 | 0.005 | -0.328 | <0.001 | -0.136 | 0.301 | -0.205 | 0.026 | -0.205 | 0.006 |
| 3 & 8 | -0.240 | 0.017 | -0.132 | 0.430 | -0.061 | 0.566 | -0.113 | 0.204 | -0.302 | 0.019 | -0.110 | 0.236 | -0.222 | 0.003 |
| 3 & 9 | -0.134 | 0.189 | 0.009 | 0.959 | -0.453 | <0.001 | -0.394 | <0.001 | -0.259 | 0.046 | -0.069 | 0.455 | -0.145 | 0.053 |
| 3 & 10 | -0.095 | 0.350 | -0.130 | 0.436 | -0.447 | <0.001 | -0.396 | <0.001 | -0.144 | 0.271 | -0.323 | <0.001 | -0.293 | <0.001 |
| 3 & 11 | -0.082 | 0.420 | -0.102 | 0.542 | -0.234 | 0.027 | -0.209 | 0.018 | -0.009 | 0.944 | -0.119 | 0.200 | -0.061 | 0.417 |
| 3 & 12 | -0.090 | 0.380 | -0.265 | 0.108 | -0.266 | 0.011 | -0.299 | 0.001 | -0.004 | 0.976 | -0.259 | 0.005 | -0.225 | 0.003 |
| 3 & 13 | -0.221 | 0.029 | -0.347 | 0.033 | -0.479 | <0.001 | -0.450 | <0.001 | -0.172 | 0.189 | -0.208 | 0.024 | -0.200 | 0.007 |
| 3 & 14 | -0.136 | 0.183 | -0.189 | 0.255 | -0.279 | 0.008 | -0.289 | 0.001 | -0.195 | 0.134 | -0.236 | 0.010 | -0.235 | 0.002 |
| 3 & 15 | -0.135 | 0.186 | -0.159 | 0.341 | -0.377 | <0.001 | -0.357 | <0.001 | -0.129 | 0.326 | -0.240 | 0.009 | -0.230 | 0.002 |
| 3 & 16 | -0.017 | 0.865 | -0.080 | 0.634 | -0.193 | 0.069 | -0.210 | 0.017 | -0.004 | 0.977 | -0.312 | 0.001 | -0.273 | <0.001 |
| 3 & 17 | -0.217 | 0.032 | -0.171 | 0.306 | -0.402 | <0.001 | -0.367 | <0.001 | -0.262 | 0.044 | -0.306 | 0.001 | -0.303 | <0.001 |
| 3 & 18 | -0.220 | 0.030 | -0.202 | 0.223 | * | * | -0.073 | 0.411 | -0.197 | 0.131 | -0.201 | 0.029 | -0.191 | 0.011 |
| 3 & 19 | -0.181 | 0.074 | 0.111 | 0.509 | 0.263 | 0.012 | 0.185 | 0.037 | -0.362 | 0.004 | 0.098 | 0.289 | -0.069 | 0.363 |
| 3 & 20 | -0.215 | 0.034 | -0.344 | 0.034 | -0.212 | 0.045 | -0.229 | 0.009 | -0.158 | 0.228 | -0.128 | 0.166 | -0.152 | 0.042 |
| 4 & 5 | 0.325 | 0.001 | 0.518 | 0.001 | 0.206 | 0.052 | 0.280 | 0.001 | 0.185 | 0.157 | 0.375 | <0.001 | 0.259 | <0.001 |
| 4 & 6 | 0.155 | 0.128 | -0.149 | 0.373 | 0.099 | 0.354 | 0.022 | 0.804 | 0.383 | 0.003 | 0.257 | 0.005 | 0.322 | <0.001 |
| 4 & 7 | -0.029 | 0.775 | 0.184 | 0.270 | 0.011 | 0.918 | 0.110 | 0.218 | -0.123 | 0.348 | -0.272 | 0.003 | -0.178 | 0.017 |
| 4 & 8 | 0.011 | 0.911 | 0.115 | 0.490 | -0.098 | 0.358 | -0.014 | 0.873 | -0.081 | 0.538 | -0.328 | <0.001 | -0.193 | 0.010 |
| 4 & 9 | 0.012 | 0.908 | 0.186 | 0.264 | -0.008 | 0.940 | 0.090 | 0.311 | -0.081 | 0.537 | -0.115 | 0.215 | -0.069 | 0.360 |
| 4 & 10 | -0.007 | 0.942 | 0.082 | 0.625 | -0.113 | 0.289 | -0.007 | 0.941 | -0.011 | 0.936 | -0.307 | 0.001 | -0.153 | 0.041 |
| 4 & 11 | 0.068 | 0.506 | 0.099 | 0.556 | -0.146 | 0.168 | -0.073 | 0.411 | 0.014 | 0.917 | 0.135 | 0.146 | 0.058 | 0.443 |
| 4 & 12 | 0.041 | 0.687 | 0.156 | 0.351 | 0.025 | 0.813 | 0.097 | 0.278 | 0.055 | 0.679 | -0.237 | 0.010 | -0.107 | 0.155 |

| | All Samples - Baseline only | | Control – Baseline only | | Control – Trial only | | Control – Baseline & Trial | | Treatment – Baseline only | | Treatment – Trial only | | Treatment – Baseline & Trial | |
|---|---|---|---|---|---|---|---|---|---|---|---|---|---|---|
| OTU set | R | *p*-value | R | *p*-value | R | *p*-value | R | *p*-value | R | *p*-value | R | *p*-value | R | *p*-value |
| 4 & 13 | -0.193 | 0.057 | 0.098 | 0.559 | -0.148 | 0.165 | -0.067 | 0.456 | -0.342 | 0.008 | -0.214 | 0.020 | -0.256 | 0.001 |
| 4 & 14 | 0.103 | 0.314 | 0.122 | 0.465 | 0.058 | 0.590 | 0.096 | 0.281 | 0.194 | 0.137 | -0.244 | 0.008 | -0.109 | 0.148 |
| 4 & 15 | 0.004 | 0.966 | 0.151 | 0.367 | 0.017 | 0.870 | 0.070 | 0.431 | -0.066 | 0.618 | -0.265 | 0.004 | -0.179 | 0.017 |
| 4 & 16 | -0.007 | 0.945 | 0.104 | 0.535 | -0.054 | 0.614 | 0.016 | 0.855 | -0.076 | 0.563 | -0.270 | 0.003 | -0.147 | 0.050 |
| 4 & 17 | -0.046 | 0.655 | 0.141 | 0.397 | -0.220 | 0.037 | -0.109 | 0.219 | -0.128 | 0.331 | -0.288 | 0.002 | -0.223 | 0.003 |
| 4 & 18 | 0.105 | 0.304 | -0.008 | 0.963 | * | * | -0.010 | 0.912 | 0.076 | 0.562 | 0.180 | 0.051 | 0.098 | 0.194 |
| 4 & 19 | 0.186 | 0.066 | 0.168 | 0.313 | 0.360 | <0.001 | 0.327 | <0.001 | 0.186 | 0.154 | -0.010 | 0.917 | 0.080 | 0.288 |
| 4 & 20 | 0.083 | 0.418 | 0.257 | 0.120 | -0.075 | 0.481 | 0.014 | 0.879 | 0.013 | 0.922 | -0.171 | 0.064 | -0.115 | 0.127 |
| 5 & 6 | 0.100 | 0.329 | -0.124 | 0.459 | 0.060 | 0.575 | 0.010 | 0.907 | 0.222 | 0.088 | 0.073 | 0.434 | 0.125 | 0.096 |
| 5 & 7 | -0.031 | 0.759 | -0.076 | 0.651 | -0.174 | 0.101 | -0.143 | 0.106 | -0.057 | 0.667 | -0.274 | 0.003 | -0.195 | 0.009 |
| 5 & 8 | -0.030 | 0.770 | -0.131 | 0.432 | -0.029 | 0.786 | -0.092 | 0.301 | 0.055 | 0.679 | -0.028 | 0.760 | 0.000 | 0.996 |
| 5 & 9 | 0.028 | 0.783 | -0.094 | 0.574 | -0.237 | 0.025 | -0.214 | 0.016 | 0.108 | 0.412 | -0.214 | 0.020 | -0.106 | 0.158 |
| 5 & 10 | 0.042 | 0.683 | -0.127 | 0.447 | -0.344 | 0.001 | -0.280 | 0.001 | 0.149 | 0.257 | -0.234 | 0.011 | -0.111 | 0.140 |
| 5 & 11 | 0.176 | 0.082 | -0.068 | 0.687 | -0.241 | 0.022 | -0.191 | 0.031 | 0.363 | 0.004 | 0.188 | 0.042 | 0.246 | 0.001 |
| 5 & 12 | 0.060 | 0.561 | -0.139 | 0.404 | -0.232 | 0.028 | -0.209 | 0.018 | 0.206 | 0.114 | -0.207 | 0.025 | -0.094 | 0.213 |
| 5 & 13 | -0.064 | 0.528 | -0.013 | 0.937 | -0.361 | <0.001 | -0.263 | 0.003 | -0.109 | 0.407 | -0.208 | 0.024 | -0.175 | 0.019 |
| 5 & 14 | 0.005 | 0.959 | -0.029 | 0.862 | -0.219 | 0.038 | -0.186 | 0.035 | -0.027 | 0.836 | -0.246 | 0.007 | -0.186 | 0.013 |
| 5 & 15 | -0.013 | 0.896 | 0.065 | 0.697 | -0.200 | 0.058 | -0.135 | 0.128 | -0.093 | 0.480 | -0.261 | 0.004 | -0.206 | 0.006 |
| 5 & 16 | 0.124 | 0.223 | 0.076 | 0.651 | -0.193 | 0.068 | -0.145 | 0.103 | 0.152 | 0.246 | -0.323 | <0.001 | -0.224 | 0.003 |
| 5 & 17 | 0.000 | 1.000 | 0.065 | 0.700 | -0.295 | 0.005 | -0.192 | 0.029 | -0.059 | 0.653 | -0.342 | <0.001 | -0.247 | 0.001 |
| 5 & 18 | 0.319 | 0.001 | 0.031 | 0.854 | * | * | 0.042 | 0.638 | 0.478 | <0.001 | 0.377 | <0.001 | 0.413 | <0.001 |
| 5 & 19 | -0.007 | 0.947 | 0.086 | 0.609 | 0.197 | 0.063 | 0.143 | 0.108 | -0.066 | 0.616 | -0.222 | 0.016 | -0.174 | 0.020 |
| 5 & 20 | 0.040 | 0.694 | 0.015 | 0.928 | -0.170 | 0.108 | -0.096 | 0.279 | 0.040 | 0.762 | -0.218 | 0.018 | -0.140 | 0.062 |
| 6 & 7 | -0.124 | 0.225 | -0.129 | 0.441 | -0.259 | 0.014 | -0.238 | 0.007 | -0.235 | 0.071 | -0.361 | <0.001 | -0.280 | <0.001 |
| 6 & 8 | -0.269 | 0.007 | -0.195 | 0.240 | -0.095 | 0.374 | -0.129 | 0.146 | -0.265 | 0.041 | -0.188 | 0.041 | -0.170 | 0.023 |
| 6 & 9 | -0.118 | 0.246 | -0.138 | 0.408 | -0.218 | 0.039 | -0.209 | 0.018 | -0.137 | 0.298 | -0.403 | <0.001 | -0.281 | <0.001 |
| 6 & 10 | -0.019 | 0.850 | -0.043 | 0.798 | -0.287 | 0.006 | -0.252 | 0.004 | -0.086 | 0.513 | -0.372 | <0.001 | -0.242 | 0.001 |
| 6 & 11 | -0.159 | 0.118 | -0.156 | 0.349 | 0.072 | 0.498 | -0.008 | 0.925 | -0.051 | 0.701 | 0.090 | 0.331 | 0.021 | 0.779 |
| 6 & 12 | -0.082 | 0.422 | -0.322 | 0.049 | -0.372 | <0.001 | -0.355 | <0.001 | -0.040 | 0.761 | -0.314 | 0.001 | -0.189 | 0.011 |
| 6 & 13 | -0.171 | 0.092 | -0.119 | 0.476 | -0.098 | 0.358 | -0.128 | 0.151 | -0.228 | 0.079 | -0.280 | 0.002 | -0.260 | <0.001 |
| 6 & 14 | -0.031 | 0.764 | -0.152 | 0.361 | -0.276 | 0.009 | -0.249 | 0.005 | 0.011 | 0.932 | -0.366 | <0.001 | -0.233 | 0.002 |
| 6 & 15 | -0.072 | 0.482 | -0.238 | 0.149 | -0.024 | 0.822 | -0.080 | 0.366 | 0.004 | 0.978 | -0.205 | 0.026 | -0.121 | 0.108 |

| | All Samples - Baseline only | | Control – Baseline only | | Control – Trial only | | Control – Baseline & Trial | | Treatment – Baseline only | | Treatment – Trial only | | Treatment – Baseline & Trial | |
|---|---|---|---|---|---|---|---|---|---|---|---|---|---|---|
| OTU set | R | *p*-value | R | *p*-value | R | *p*-value | R | *p*-value | R | *p*-value | R | *p*-value | R | *p*-value |
| 6 & 16 | -0.090 | 0.379 | -0.215 | 0.195 | -0.255 | 0.015 | -0.254 | 0.004 | -0.060 | 0.648 | -0.360 | <0.001 | -0.229 | 0.002 |
| 6 & 17 | -0.267 | 0.008 | -0.317 | 0.053 | -0.091 | 0.394 | -0.165 | 0.062 | -0.270 | 0.037 | -0.244 | 0.008 | -0.249 | 0.001 |
| 6 & 18 | -0.093 | 0.364 | -0.247 | 0.134 | * | * | -0.120 | 0.177 | 0.010 | 0.937 | 0.014 | 0.877 | 0.010 | 0.895 |
| 6 & 19 | -0.066 | 0.519 | -0.171 | 0.304 | -0.029 | 0.789 | -0.069 | 0.440 | 0.002 | 0.989 | -0.226 | 0.014 | -0.137 | 0.068 |
| 6 & 20 | -0.162 | 0.111 | -0.354 | 0.029 | -0.132 | 0.216 | -0.199 | 0.024 | -0.079 | 0.546 | -0.091 | 0.328 | -0.079 | 0.292 |
| 7 & 8 | 0.132 | 0.196 | 0.210 | 0.206 | 0.103 | 0.332 | 0.206 | 0.020 | 0.081 | 0.540 | 0.099 | 0.286 | 0.176 | 0.019 |
| 7 & 9 | 0.476 | <0.001 | 0.649 | <0.001 | 0.814 | <0.001 | 0.810 | <0.001 | 0.331 | 0.010 | 0.719 | <0.001 | 0.654 | <0.001 |
| 7 & 10 | 0.558 | <0.001 | 0.650 | <0.001 | 0.827 | <0.001 | 0.809 | <0.001 | 0.465 | <0.001 | 0.677 | <0.001 | 0.642 | <0.001 |
| 7 & 11 | 0.209 | 0.039 | 0.600 | <0.001 | 0.252 | 0.017 | 0.329 | <0.001 | 0.027 | 0.837 | -0.011 | 0.907 | -0.014 | 0.854 |
| 7 & 12 | 0.666 | <0.001 | 0.803 | <0.001 | 0.772 | <0.001 | 0.781 | <0.001 | 0.489 | <0.001 | 0.612 | <0.001 | 0.587 | <0.001 |
| 7 & 13 | 0.312 | 0.002 | 0.435 | 0.006 | 0.547 | <0.001 | 0.507 | <0.001 | 0.201 | 0.124 | 0.500 | <0.001 | 0.407 | <0.001 |
| 7 & 14 | 0.222 | 0.028 | 0.346 | 0.033 | 0.536 | <0.001 | 0.520 | <0.001 | -0.065 | 0.624 | 0.644 | <0.001 | 0.558 | <0.001 |
| 7 & 15 | 0.169 | 0.096 | 0.391 | 0.015 | 0.435 | <0.001 | 0.431 | <0.001 | -0.066 | 0.617 | 0.480 | <0.001 | 0.389 | <0.001 |
| 7 & 16 | 0.347 | <0.001 | 0.586 | <0.001 | 0.603 | <0.001 | 0.612 | <0.001 | -0.065 | 0.624 | 0.660 | <0.001 | 0.575 | <0.001 |
| 7 & 17 | 0.290 | 0.004 | 0.513 | 0.001 | 0.564 | <0.001 | 0.554 | <0.001 | 0.103 | 0.434 | 0.396 | <0.001 | 0.317 | <0.001 |
| 7 & 18 | -0.035 | 0.730 | 0.196 | 0.239 | * | * | 0.016 | 0.858 | -0.016 | 0.906 | -0.162 | 0.080 | -0.122 | 0.105 |
| 7 & 19 | 0.152 | 0.134 | 0.328 | 0.045 | -0.104 | 0.331 | 0.059 | 0.508 | 0.045 | 0.735 | 0.082 | 0.378 | 0.088 | 0.243 |
| 7 & 20 | 0.375 | <0.001 | 0.491 | 0.002 | 0.283 | 0.007 | 0.277 | 0.002 | 0.232 | 0.075 | 0.305 | 0.001 | 0.286 | <0.001 |
| 8 & 9 | 0.228 | 0.024 | 0.231 | 0.164 | 0.133 | 0.212 | 0.218 | 0.013 | 0.239 | 0.066 | 0.228 | 0.013 | 0.303 | <0.001 |
| 8 & 10 | 0.126 | 0.215 | 0.122 | 0.465 | 0.126 | 0.238 | 0.178 | 0.044 | 0.177 | 0.177 | 0.171 | 0.064 | 0.271 | <0.001 |
| 8 & 11 | 0.284 | 0.005 | 0.417 | 0.009 | 0.025 | 0.814 | 0.142 | 0.111 | 0.201 | 0.124 | 0.016 | 0.863 | 0.048 | 0.525 |
| 8 & 12 | 0.068 | 0.505 | 0.118 | 0.480 | 0.097 | 0.365 | 0.145 | 0.101 | 0.050 | 0.703 | 0.143 | 0.123 | 0.210 | 0.005 |
| 8 & 13 | 0.066 | 0.519 | 0.213 | 0.199 | -0.045 | 0.670 | 0.047 | 0.600 | -0.037 | 0.777 | 0.180 | 0.052 | 0.135 | 0.072 |
| 8 & 14 | 0.125 | 0.220 | 0.143 | 0.392 | 0.101 | 0.345 | 0.153 | 0.084 | 0.146 | 0.265 | 0.063 | 0.500 | 0.155 | 0.039 |
| 8 & 15 | 0.091 | 0.373 | 0.157 | 0.348 | -0.017 | 0.873 | 0.067 | 0.450 | 0.044 | 0.741 | 0.205 | 0.026 | 0.218 | 0.004 |
| 8 & 16 | 0.026 | 0.803 | -0.063 | 0.706 | 0.072 | 0.498 | 0.103 | 0.246 | 0.211 | 0.106 | 0.091 | 0.324 | 0.209 | 0.005 |
| 8 & 17 | 0.141 | 0.166 | 0.255 | 0.123 | -0.094 | 0.380 | 0.022 | 0.809 | 0.050 | 0.706 | 0.141 | 0.129 | 0.143 | 0.056 |
| 8 & 18 | 0.251 | 0.013 | -0.103 | 0.538 | * | * | -0.080 | 0.368 | 0.357 | 0.005 | 0.036 | 0.701 | 0.120 | 0.110 |
| 8 & 19 | 0.166 | 0.102 | 0.298 | 0.069 | -0.039 | 0.715 | 0.078 | 0.379 | 0.073 | 0.578 | 0.161 | 0.081 | 0.153 | 0.042 |
| 8 & 20 | 0.176 | 0.084 | 0.147 | 0.377 | 0.191 | 0.071 | 0.174 | 0.049 | 0.215 | 0.099 | 0.080 | 0.388 | 0.140 | 0.063 |
| 9 & 10 | 0.613 | <0.001 | 0.586 | <0.001 | 0.866 | <0.001 | 0.839 | <0.001 | 0.669 | <0.001 | 0.558 | <0.001 | 0.605 | <0.001 |
| 9 & 11 | 0.332 | 0.001 | 0.436 | 0.006 | 0.374 | <0.001 | 0.394 | <0.001 | 0.289 | 0.025 | -0.032 | 0.729 | 0.033 | 0.667 |

| | All Samples - Baseline only | | Control – Baseline only | | Control – Trial only | | Control – Baseline & Trial | | Treatment – Baseline only | | Treatment – Trial only | | Treatment – Baseline & Trial | |
|---|---|---|---|---|---|---|---|---|---|---|---|---|---|---|
| OTU set | R | *p*-value | R | *p*-value | R | *p*-value | R | *p*-value | R | *p*-value | R | *p*-value | R | *p*-value |
| 9 & 12 | 0.380 | <0.001 | 0.500 | 0.001 | 0.764 | <0.001 | 0.736 | <0.001 | 0.266 | 0.040 | 0.567 | <0.001 | 0.530 | <0.001 |
| 9 & 13 | 0.220 | 0.029 | 0.218 | 0.188 | 0.558 | <0.001 | 0.501 | <0.001 | 0.226 | 0.082 | 0.628 | <0.001 | 0.502 | <0.001 |
| 9 & 14 | 0.232 | 0.022 | 0.261 | 0.113 | 0.535 | <0.001 | 0.511 | <0.001 | 0.210 | 0.108 | 0.545 | <0.001 | 0.520 | <0.001 |
| 9 & 15 | 0.122 | 0.230 | 0.212 | 0.201 | 0.369 | <0.001 | 0.357 | <0.001 | 0.048 | 0.714 | 0.414 | <0.001 | 0.363 | <0.001 |
| 9 & 16 | 0.269 | 0.007 | 0.501 | 0.001 | 0.545 | <0.001 | 0.571 | <0.001 | -0.065 | 0.624 | 0.564 | <0.001 | 0.520 | <0.001 |
| 9 & 17 | 0.246 | 0.015 | 0.412 | 0.010 | 0.527 | <0.001 | 0.517 | <0.001 | 0.125 | 0.340 | 0.328 | <0.001 | 0.272 | <0.001 |
| 9 & 18 | -0.046 | 0.654 | -0.090 | 0.589 | * | * | -0.090 | 0.311 | -0.023 | 0.863 | -0.144 | 0.119 | -0.111 | 0.142 |
| 9 & 19 | 0.378 | <0.001 | 0.539 | <0.001 | -0.183 | 0.083 | 0.044 | 0.619 | 0.265 | 0.041 | 0.193 | 0.037 | 0.216 | 0.004 |
| 9 & 20 | 0.170 | 0.095 | 0.178 | 0.284 | 0.308 | 0.003 | 0.231 | 0.009 | 0.156 | 0.233 | 0.400 | <0.001 | 0.346 | <0.001 |
| 10 & 11 | 0.370 | <0.001 | 0.668 | <0.001 | 0.337 | 0.001 | 0.418 | <0.001 | 0.272 | 0.036 | 0.228 | 0.013 | 0.223 | 0.003 |
| 10 & 12 | 0.447 | <0.001 | 0.559 | <0.001 | 0.807 | <0.001 | 0.783 | <0.001 | 0.296 | 0.021 | 0.756 | <0.001 | 0.677 | <0.001 |
| 10 & 13 | 0.341 | 0.001 | 0.429 | 0.007 | 0.623 | <0.001 | 0.583 | <0.001 | 0.259 | 0.046 | 0.674 | <0.001 | 0.537 | <0.001 |
| 10 & 14 | 0.334 | 0.001 | 0.365 | 0.024 | 0.585 | <0.001 | 0.563 | <0.001 | 0.279 | 0.031 | 0.613 | <0.001 | 0.566 | <0.001 |
| 10 & 15 | 0.238 | 0.018 | 0.327 | 0.045 | 0.408 | <0.001 | 0.411 | <0.001 | 0.155 | 0.236 | 0.551 | <0.001 | 0.479 | <0.001 |
| 10 & 16 | 0.357 | <0.001 | 0.596 | <0.001 | 0.612 | <0.001 | 0.635 | <0.001 | -0.061 | 0.642 | 0.773 | <0.001 | 0.676 | <0.001 |
| 10 & 17 | 0.256 | 0.011 | 0.434 | 0.007 | 0.602 | <0.001 | 0.583 | <0.001 | 0.117 | 0.373 | 0.609 | <0.001 | 0.469 | <0.001 |
| 10 & 18 | -0.080 | 0.436 | -0.128 | 0.445 | * | * | -0.096 | 0.283 | 0.002 | 0.990 | -0.032 | 0.731 | -0.019 | 0.801 |
| 10 & 19 | 0.206 | 0.042 | 0.146 | 0.381 | -0.263 | 0.012 | -0.094 | 0.289 | 0.293 | 0.023 | 0.092 | 0.320 | 0.159 | 0.034 |
| 10 & 20 | 0.287 | 0.004 | 0.381 | 0.018 | 0.342 | 0.001 | 0.331 | <0.001 | 0.170 | 0.195 | 0.448 | <0.001 | 0.363 | <0.001 |
| 11 & 12 | 0.263 | 0.009 | 0.402 | 0.012 | 0.286 | 0.006 | 0.314 | <0.001 | 0.277 | 0.032 | 0.022 | 0.813 | 0.055 | 0.463 |
| 11 & 13 | 0.265 | 0.008 | 0.449 | 0.005 | 0.316 | 0.002 | 0.368 | <0.001 | 0.190 | 0.146 | 0.110 | 0.235 | 0.132 | 0.079 |
| 11 & 14 | 0.170 | 0.094 | 0.328 | 0.044 | 0.151 | 0.156 | 0.185 | 0.037 | 0.074 | 0.574 | -0.118 | 0.203 | -0.101 | 0.180 |
| 11 & 15 | 0.108 | 0.288 | 0.237 | 0.153 | 0.076 | 0.475 | 0.134 | 0.131 | 0.042 | 0.752 | 0.074 | 0.426 | 0.044 | 0.561 |
| 11 & 16 | 0.318 | 0.001 | 0.555 | <0.001 | 0.207 | 0.051 | 0.276 | 0.002 | 0.156 | 0.235 | 0.003 | 0.975 | 0.004 | 0.955 |
| 11 & 17 | 0.189 | 0.062 | 0.464 | 0.003 | 0.286 | 0.006 | 0.342 | <0.001 | 0.037 | 0.777 | 0.120 | 0.196 | 0.090 | 0.231 |
| 11 & 18 | 0.246 | 0.015 | -0.127 | 0.446 | * | * | -0.075 | 0.399 | 0.295 | 0.022 | 0.188 | 0.041 | 0.231 | 0.002 |
| 11 & 19 | 0.190 | 0.061 | 0.250 | 0.131 | -0.019 | 0.859 | 0.060 | 0.503 | 0.152 | 0.246 | -0.047 | 0.610 | 0.007 | 0.924 |
| 11 & 20 | 0.251 | 0.013 | 0.439 | 0.006 | 0.150 | 0.158 | 0.244 | 0.006 | 0.155 | 0.237 | 0.111 | 0.231 | 0.115 | 0.125 |
| 12 & 13 | 0.272 | 0.007 | 0.254 | 0.124 | 0.425 | <0.001 | 0.400 | <0.001 | 0.263 | 0.042 | 0.710 | <0.001 | 0.578 | <0.001 |
| 12 & 14 | 0.295 | 0.003 | 0.396 | 0.014 | 0.628 | <0.001 | 0.591 | <0.001 | -0.043 | 0.742 | 0.680 | <0.001 | 0.626 | <0.001 |
| 12 & 15 | 0.159 | 0.117 | 0.416 | 0.009 | 0.257 | 0.014 | 0.318 | <0.001 | -0.139 | 0.289 | 0.549 | <0.001 | 0.451 | <0.001 |
| 12 & 16 | 0.394 | <0.001 | 0.541 | <0.001 | 0.618 | <0.001 | 0.612 | <0.001 | -0.043 | 0.742 | 0.742 | <0.001 | 0.680 | <0.001 |

| | All Samples - Baseline only | | Control – Baseline only | | Control – Trial only | | Control – Baseline & Trial | | Treatment – Baseline only | | Treatment – Trial only | | Treatment – Baseline & Trial | |
|---|---|---|---|---|---|---|---|---|---|---|---|---|---|---|
| OTU set | R | *p*-value | R | *p*-value | R | *p*-value | R | *p*-value | R | *p*-value | R | *p*-value | R | *p*-value |
| 12 & 17 | 0.240 | 0.017 | 0.371 | 0.022 | 0.390 | <0.001 | 0.405 | <0.001 | 0.078 | 0.554 | 0.577 | <0.001 | 0.454 | <0.001 |
| 12 & 18 | 0.069 | 0.497 | 0.246 | 0.136 | * | * | 0.053 | 0.551 | 0.168 | 0.199 | -0.025 | 0.791 | 0.020 | 0.791 |
| 12 & 19 | 0.128 | 0.210 | 0.163 | 0.328 | -0.184 | 0.082 | -0.051 | 0.567 | 0.139 | 0.289 | 0.057 | 0.538 | 0.088 | 0.241 |
| 12 & 20 | 0.457 | <0.001 | 0.428 | 0.007 | 0.307 | 0.003 | 0.301 | 0.001 | 0.485 | <0.001 | 0.477 | <0.001 | 0.466 | <0.001 |
| 13 & 14 | 0.030 | 0.766 | 0.073 | 0.665 | 0.508 | <0.001 | 0.431 | <0.001 | -0.083 | 0.526 | 0.529 | <0.001 | 0.408 | <0.001 |
| 13 & 15 | 0.516 | <0.001 | 0.537 | 0.001 | 0.611 | <0.001 | 0.600 | <0.001 | 0.487 | <0.001 | 0.573 | <0.001 | 0.550 | <0.001 |
| 13 & 16 | 0.241 | 0.017 | 0.364 | 0.025 | 0.508 | <0.001 | 0.494 | <0.001 | 0.125 | 0.340 | 0.586 | <0.001 | 0.464 | <0.001 |
| 13 & 17 | 0.654 | <0.001 | 0.666 | <0.001 | 0.794 | <0.001 | 0.772 | <0.001 | 0.648 | <0.001 | 0.689 | <0.001 | 0.682 | <0.001 |
| 13 & 18 | -0.095 | 0.350 | 0.293 | 0.075 | * | * | 0.151 | 0.089 | -0.167 | 0.202 | -0.121 | 0.192 | -0.137 | 0.069 |
| 13 & 19 | 0.101 | 0.322 | 0.207 | 0.213 | -0.309 | 0.003 | -0.155 | 0.080 | 0.040 | 0.762 | 0.135 | 0.144 | 0.114 | 0.131 |
| 13 & 20 | 0.653 | <0.001 | 0.781 | <0.001 | 0.415 | <0.001 | 0.513 | <0.001 | 0.555 | <0.001 | 0.595 | <0.001 | 0.586 | <0.001 |
| 14 & 15 | 0.162 | 0.112 | 0.291 | 0.076 | 0.216 | 0.040 | 0.251 | 0.004 | -0.054 | 0.679 | 0.432 | <0.001 | 0.396 | <0.001 |
| 14 & 16 | 0.378 | <0.001 | 0.447 | 0.005 | 0.555 | <0.001 | 0.540 | <0.001 | -0.017 | 0.898 | 0.763 | <0.001 | 0.739 | <0.001 |
| 14 & 17 | 0.146 | 0.151 | 0.120 | 0.472 | 0.373 | <0.001 | 0.339 | <0.001 | 0.193 | 0.140 | 0.348 | <0.001 | 0.303 | <0.001 |
| 14 & 18 | -0.086 | 0.398 | -0.056 | 0.737 | * | * | -0.050 | 0.579 | -0.061 | 0.642 | -0.171 | 0.064 | -0.142 | 0.059 |
| 14 & 19 | 0.249 | 0.013 | 0.301 | 0.066 | -0.231 | 0.029 | -0.091 | 0.308 | 0.237 | 0.069 | 0.177 | 0.056 | 0.185 | 0.014 |
| 14 & 20 | 0.095 | 0.351 | 0.171 | 0.304 | 0.325 | 0.002 | 0.264 | 0.003 | -0.071 | 0.590 | 0.395 | <0.001 | 0.320 | <0.001 |
| 15 & 16 | 0.146 | 0.152 | 0.271 | 0.099 | 0.309 | 0.003 | 0.327 | <0.001 | -0.054 | 0.679 | 0.472 | <0.001 | 0.430 | <0.001 |
| 15 & 17 | 0.382 | <0.001 | 0.544 | <0.001 | 0.454 | <0.001 | 0.482 | <0.001 | 0.261 | 0.044 | 0.615 | <0.001 | 0.528 | <0.001 |
| 15 & 18 | -0.021 | 0.841 | 0.337 | 0.039 | * | * | 0.149 | 0.093 | -0.089 | 0.497 | -0.114 | 0.220 | -0.108 | 0.152 |
| 15 & 19 | 0.121 | 0.235 | 0.276 | 0.094 | -0.031 | 0.771 | 0.057 | 0.526 | 0.023 | 0.859 | 0.073 | 0.432 | 0.077 | 0.307 |
| 15 & 20 | 0.577 | <0.001 | 0.681 | <0.001 | 0.320 | 0.002 | 0.412 | <0.001 | 0.479 | <0.001 | 0.648 | <0.001 | 0.614 | <0.001 |
| 16 & 17 | 0.373 | <0.001 | 0.521 | 0.001 | 0.474 | <0.001 | 0.495 | <0.001 | 0.218 | 0.095 | 0.515 | <0.001 | 0.430 | <0.001 |
| 16 & 18 | 0.038 | 0.712 | -0.056 | 0.737 | * | * | -0.056 | 0.534 | 0.217 | 0.095 | -0.234 | 0.011 | -0.161 | 0.031 |
| 16 & 19 | 0.152 | 0.136 | 0.332 | 0.042 | -0.177 | 0.095 | -0.030 | 0.737 | -0.065 | 0.624 | 0.145 | 0.116 | 0.136 | 0.070 |
| 16 & 20 | 0.201 | 0.047 | 0.349 | 0.032 | 0.284 | 0.007 | 0.266 | 0.002 | -0.071 | 0.590 | 0.439 | <0.001 | 0.353 | <0.001 |
| 17 & 18 | -0.047 | 0.647 | 0.310 | 0.058 | * | * | 0.153 | 0.086 | -0.119 | 0.364 | -0.130 | 0.161 | -0.126 | 0.095 |
| 17 & 19 | 0.230 | 0.022 | 0.501 | 0.001 | -0.312 | 0.003 | -0.099 | 0.266 | 0.054 | 0.682 | 0.024 | 0.797 | 0.039 | 0.603 |
| 17 & 20 | 0.532 | <0.001 | 0.690 | <0.001 | 0.341 | 0.001 | 0.430 | <0.001 | 0.402 | 0.001 | 0.510 | <0.001 | 0.484 | <0.001 |
| 18 & 19 | 0.027 | 0.794 | 0.277 | 0.092 | * | * | 0.071 | 0.426 | -0.030 | 0.818 | -0.160 | 0.084 | -0.121 | 0.109 |
| 18 & 20 | 0.055 | 0.591 | 0.318 | 0.052 | * | * | 0.180 | 0.042 | 0.056 | 0.669 | -0.166 | 0.072 | -0.102 | 0.174 |
| 19 & 20 | 0.069 | 0.497 | 0.156 | 0.350 | -0.143 | 0.180 | -0.076 | 0.396 | 0.030 | 0.819 | 0.119 | 0.201 | 0.103 | 0.170 |

* - No OTU18 were present in Control animals during the Trial period.

**S2 Table. Change in OTU relative abundance from baseline to treatment periods for Control animals.**

| | | All samples | | Proestrus | | Estrus | | Metestrus | | Diestrus | |
|---|---|---|---|---|---|---|---|---|---|---|---|
| | Genus | B:T Ratio | *p*-value | B:T Ratio | *p*-value | B:T Ratio | *p*-value | B:T Ratio | *p*-value | B:T Ratio | *p*-value |
| OTU1 | *Proteus* | 0.913 | 0.491 | 1.000 | 1.000 | 0.952 | 0.852 | 0.758 | 0.315 | 0.963 | 0.912 |
| OTU2 | *Escherichia/Shigella* | 1.342 | 0.027 | 2.100 | 0.030 | 1.403 | 0.152 | 1.500 | 0.156 | 0.850 | 0.529 |
| OTU3 | *Streptococcus* | 1.313 | 0.041 | 0.732 | 0.429 | 1.350 | 0.205 | 1.704 | 0.043 | 1.295 | 0.315 |
| OTU4 | *Morganella* | 0.857 | 0.227 | 0.885 | 0.792 | 0.832 | 0.437 | 0.919 | 0.780 | 0.765 | 0.315 |
| OTU5 | *Pasteurellaceae* | 1.119 | 0.396 | 1.848 | 0.082 | 0.801 | 0.347 | 1.344 | 0.278 | 1.132 | 0.631 |
| OTU6 | *Enterococcus* | 1.036 | 0.791 | 1.063 | 0.931 | 0.994 | 0.979 | 0.979 | 0.968 | 1.199 | 0.481 |
| OTU7 | *Corynebacterium* | 1.622 | < 0.001 | 0.805 | 0.537 | 0.577 | 0.016 | 0.488 | 0.008 | 0.641 | 0.089 |

**S3 Table. Change in OTU relative abundance from baseline to treatment periods for Treatment animals.**

| | | All samples | | Proestrus | | Estrus | | Metestrus | | Diestrus | |
|---|---|---|---|---|---|---|---|---|---|---|---|
| | Genus | B:T Ratio | *p*-value | B:T Ratio | *p*-value | B:T Ratio | *p*-value | B:T Ratio | *p*-value | B:T Ratio | *p*-value |
| OTU1 | *Proteus* | 1.066 | 0.560 | 1.375 | 0.371 | 1.308 | 0.241 | 0.772 | 0.368 | 1.068 | 0.699 |
| OTU2 | *Escherichia/Shigella* | 0.997 | 0.976 | 1.214 | 0.594 | 0.703 | 0.100 | 0.788 | 0.407 | 1.151 | 0.413 |
| OTU3 | *Streptococcus* | 1.236 | 0.056 | 1.058 | 0.859 | 1.695 | 0.020 | 1.123 | 0.680 | 0.920 | 0.624 |
| OTU4 | *Morganella* | 0.813 | 0.057 | 1.318 | 0.440 | 0.914 | 0.698 | 0.725 | 0.267 | 0.650 | 0.011 |
| OTU5 | *Pasteurellaceae* | 1.027 | 0.799 | 0.963 | 0.953 | 0.854 | 0.478 | 1.152 | 0.581 | 1.198 | 0.273 |
| OTU6 | *Enterococcus* | 0.831 | 0.090 | 0.318 | 0.440 | 0.905 | 0.664 | 0.756 | 0.332 | 0.799 | 0.187 |
| OTU7 | *Corynebacterium* | 1.187 | 0.049 | 0.517 | 0.019 | 0.788 | 0.280 | 0.877 | 0.630 | 0.895 | 0.481 |
